# Depicting pathogenesis of osteomyelitis by single cell RNA-sequencing and an involvement of *Morrbid* in the autoinflammatory disease

**DOI:** 10.1101/2025.01.23.634426

**Authors:** Qingran Huo, Jiayu Ding, Hongxi Zhou, Hang He, Lorie Chen Cai, Jingjing Liu, Ge Dong, Zhigang Cai

## Abstract

Autoinflammatory diseases (AIDs) are defined as abnormal activation of the innate immune system leading to spontaneous and uncontrolled inflammation. The AIDs affect bone tissue and lead to chronic recurrent multifocal osteomyelitis (CRMO). However, the etiology and treatment of CRMO remain elusive. A mouse strain, *Pstpip2^cmo/cmo^* (*cmo*: *chronic multifocal osteomyelitis*), exhibits phenotypic characteristics similar to human CRMO. *Morrbid* is a long non-coding RNA gene and has been indicated in leukemogenesis in our previous studies. In this study, we demonstrated that *Morrbid* and *Pstpip2* are co-expressed in mature myeloid cells and hypothesized a role of *Morrbid* in osteomyelitis. The *Pstpip2^−/−^* mice have the same phenotype as *Pstpip2^cmo/cmo^,* mimicking CRMO, while loss of *Morrbid* in *Pstpip2^−/−^*mice significantly inhibited the initiation and progression of CRMO symptoms, as well as the dysregulated activation of myeloid cells and the excessive release of inflammatory cytokines. Furthermore, single-cell RNA-sequencing (scRNA-seq) analysis from the *Pstpip2^−/−^* mice and the compound mutant mice supports that reduction of osteoclasts and inflammatory cells caused by *Morrbid* loss. The study systematically profiles the etiology of CRMO by scRNA-seq and warrants that inhibiting the lifespan of inflammatory myeloid cells by targeting *Morrbid* is an effective therapeutic strategy for osteomyelitis.

**Highlights:**

1. We generated a frameshift mutant mouse strain *Pstpip2^−/−^*, which have a classic CRMO-like phenotype as same as *Pstpip2^cmo/cmo^* and could be used for testing various anti-inflammation perturbations.
2. Loss of *Morrbid* significantly inhibited the autoinflammatory symptoms in *Pstpip2^−/−^* mice, suggesting that *Morrbid* is a novel target for mitigating CRMO.
3. scRNA-seq analysis of the affected bone marrow cells in *Pstpip2^−/−^*revealed the abnormalities of osteoclasts (OC), neutrophils (NE) and granulocyte macrophage progenitors (GMP) in both their fractions and inflammatory activities.
4. Upon loss of *Morrbid*, we observed reduced composition and proliferation of OC and decreased activity of *Nfkb2* and *Rela* in the compound mutants.

## Introduction

Inflammatory diseases caused by the instinct dysfunction of the innate immune cells are called autoinflammatory diseases (AIDs)^[1, 2]^. At present, many types of AIDs have been identified in clinic, including colitis, dermatosis, chondritis, osteomyelitis, vasculitis, and a recently reported rheumatoid-hematological VEXAS^[3–5]^. However, treatments for AIDs are limited^[4, 6]^. It is urgent to have more insightful understanding the pathogenesis of AIDs and develop more accurate and effective treatment methods.

Chronic recurrent multifocal osteomyelitis (CRMO) is a non-bacterial autoinflammatory bone disease characterized by multifocal and multiple recurrences^[7]^. The typical symptom of CRMO is pain at the bony injury site with or without swelling, which tends to occurs around the metaphysis of femur, tibia and clavicle^[7, 8]^. Patients with CRMO are frequently accompanied by other chronic inflammatory diseases or autoimmune diseases, including arthritis, psoriasis and inflammatory bowel disease (IBD)^[7, 9]^. Recently a mutation in *IL1R1* was reported to cause CRMO in a juvenile female with profound inflammatory myeloid cells in peripheral blood ^[10]^. However, only her peripheral mononuclear blood cells (PBMCs) were profiled by single-cell RNA-sequencing (scRNA-seq). Profiling the immune and stromal cells in the affected region will assist a direct understanding of the pathogenesis of CRMO.

A mouse model *Pstpip2^cmo/cmo^* (*cmo*: *chronic multifocal osteomyelitis*) shows disease characteristics similar to human CRMO and is widely used as an animal model for pathogenesis of AIDs. *Pstpip2^cmo/cmo^* mutant mice spontaneously develop CRMO starting at the 2-month-old age, typically characterized by progressive hind paw swelling and deformity, skin and ear inflammation and inflammatory bone resorption. It is believed that such inflammation is mediated by biased myelogenesis and inflammatory myeloid cells (such as neutrophils and macrophages) of the innate immune system^[11–14]^.

As an adaptor protein, PSTPIP2 (proline-serine-threonine-phosphatase-interacting protein 2) plays a role in regulating membrane curvature and maintaining cytoskeleton stability by binding to its interaction partners, and is highly correlated with the migration and activation of macrophages and osteoclasts (OC, bone-absorbing)^[15, 16]^. Current studies indicate that inflammasome, cysteine aspartate-specific protease (Caspase), cell stress and reactive oxygen species (ROS) are all involved in the pathological development for *Pstpip2^cmo/cmo^* ^[14, 17]^. Interestingly, mutations in *PSTPIP1* results in PSTPIP1-associated myeloid-related proteinemia inflammatory syndrome (PAMI), a rare autoinflammatory disease^[18]^. Nonetheless, mechanisms by which PSTPIP2 regulate myeloid cells especially the osteoclast cells have not been fully elucidated, and the cell populations, cytokines and signaling pathways involved in the occurrence of CRMO for *Pstpip2^cmo/cmo^* are also controversial. Furthermore, effective perturbations or drugs to inhibit the progression of the disease still remains to be explored.

Our previous studies have demonstrated that the conserved lncRNA *Morrbid* (myeloid RNA regulator of Bim-induced death) was specifically expressed in mature eosinophils, neutrophils (NE) and classical monocytes (Mono) and promotes the myeloid-linage leukemogenesis^[19]^. In addition, *Morrbid* was discovered to have an intimate connection with the release of various cytokines and the activity of cell signaling pathways^[20–23]^. This prompted us to ask whether *Morrbid* has a role in inhibiting the progression of AIDs including CRMO.

In addition, scRNA-seq technologies provide powerful assets as the traditional approaches for pathogenesis of immune diseases. It is believed that single-cell omics are revolutionizing the study of immunology and rheumatology^[24]^. Even at the most basic level, single-cell sequencing technologies are able to provide a systematic map (a cell atlas) for affected regions or organs and their microenvironment alteration. As mentioned above, even though a PBMC profile has been described in a CRMO patient^[10]^, so far, an extensive understanding of CRMO in the affected tissues in human or even in mice (i.e. in *Pstpip2^cmo/cmo^*) is still lacking.

In the present study, we showed that *Pstpip2* and *Morrbid* were co-expressed in myeloid cells. We generated a new *Pstpip2* frameshift mutant mouse strain *Pstpip2^−/−^*and observed its phenotype similar to classical *Pstpip2^cmo/cmo^*. To access the involvement of *Morrbid*, we generated double knock-out mutant *Pstpip2^−/−^; Morbbid^−/−^*(DKO). The deficiency of *Morrbid* in the DKO mice inhibited the progression of inflammatory symptoms and bone damage, which is very likely mediated by the decrease of myeloid cells and cytokines in the lesion site. The mitigation of CRMO in the DKO were also validated by the scRNA-seq analysis using *Pstpip2^−/−^* and DKO bone marrow cells.

## Results

### A 5-bp deletion in Pstpip2 gene leads to chronic osteomyelitis characterized by swelling of the hind paw and bone damage

Using the scRNA-seq dataset, we confirmed that *Pstpip2* was highly expressed in murine myeloid cells, especially the monocytes in wild type (WT) (Fig. 1A). We constructed a mouse strain carrying a novel frameshift mutation (*Pstpip2^−/−^*) by knocking out 5-bp in the *Pstpip2* exon 2 protein-coding region (Fig. 1B). The main lesion of osteomyelitis is located in the bilateral hind paws of mice. Continuously, we observe the hind paws of both WT and *Pstpip2^−/−^* mice of the same age, rigorously documenting the occurrence and progression of the disease. We established a scoring system: each swollen toe counts 1 point, with a total score of 10 points. Through continuous observation and scoring, we found that the hind paw swelling of *Pstpip2^−/−^* mice was characterized by progressive aggravation (Fig. 1C). Almost all *Pstpip2^−/−^* mice showed marked swelling in all toes of bilateral hind paws by 12 weeks of age (Fig. 1C), and their appearance was dramatically different from that of WT mice (Fig. 1D). We isolated popliteal lymph nodes (popLN) from 20-week-old WT and *Pstpip2^−/−^*mice and observed massive lymphomegaly of bilateral popliteal lymph nodes in *Pstpip2^−/−^* mice, suggestive of inflammatory symptoms in nearby tissues (Fig. 1E). These results confirm that the newly generated *Pstpip2^−/−^* mice cause a spontaneous, non-bacterial inflammatory disease similar to the classic *Pstpip2^cmo/cmo^* phenotype.

**Figure 1.**
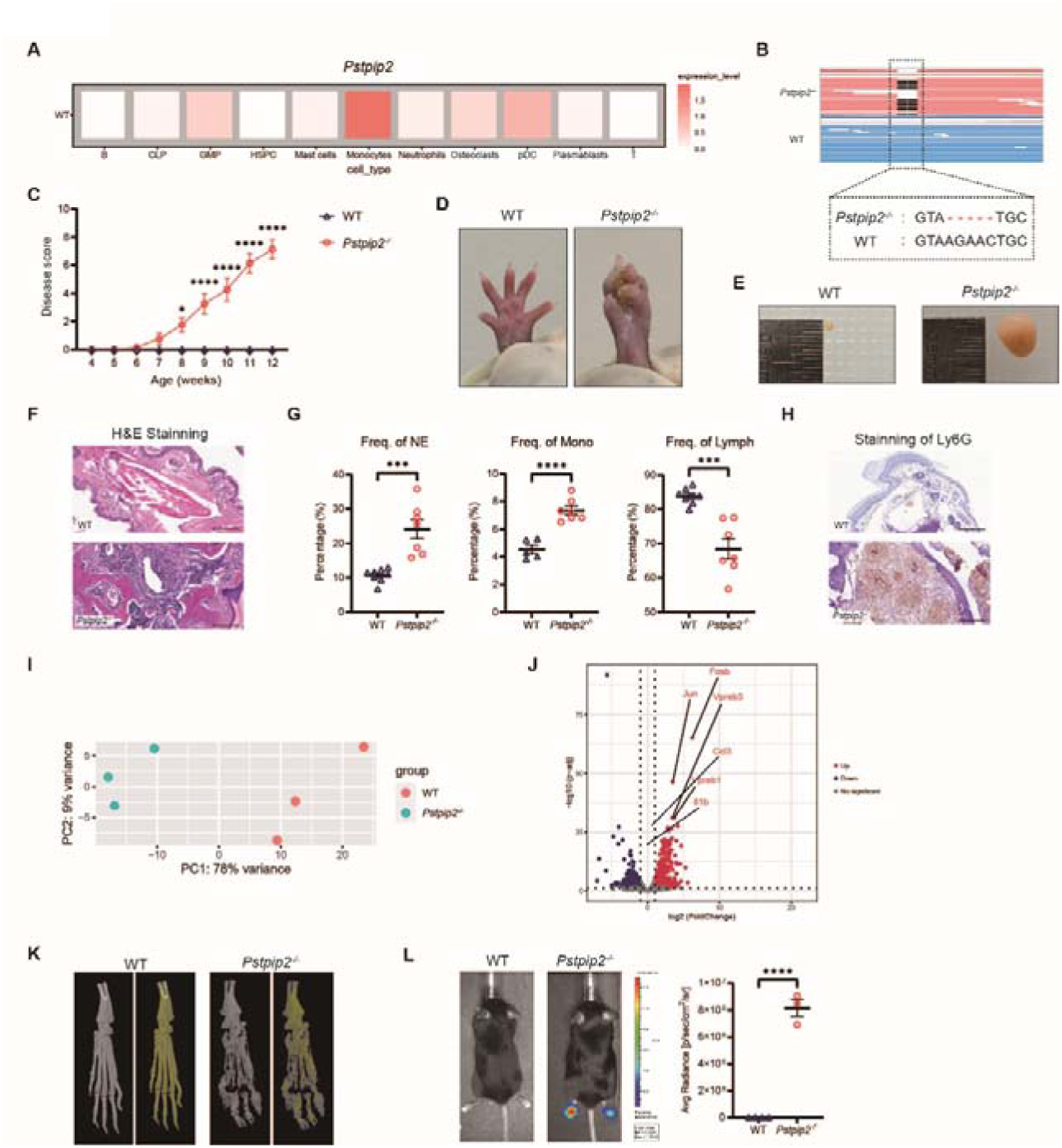
A 5-bp deletion in *Pstpip2* leads to chronic multifocal osteomyelitis in mice. (**A**) Expression of Pstpip2 in bone marrow cells (BM) revealed by scRNA-seq. See also Fig. 3 for details. (**B**) Generation of *Pstpip2^−/−^* mice (5-bp deletion in the coding region of the locus, exon 2). The upper panel shows the transcripts by the bulk RNA-seq; the lower panel shows the Sanger sequencing results. The frameshift mutation caused by the 5-bp deletion in *Pstpip2^−/−^* mice is highlighted. (**C**) Disease progression was scored (scale from 0 to 10) by visual inspection of the hind paws of WT and *Pstpip2^−/−^* mice. (n=8 per group; two-way ANOVA, mean±SEM). (**D-E)** Representative images of the hind paws and popliteal lymph nodes (popLN, e) of 20-week-old WT and *Pstpip2^−/−^* mice. (**F**) Representative hematoxylin and eosin (H&E) staining images of hind paw sections of WT and *Pstpip2^−/−^* mice (scale bar, 200 μm). (**G**) Frequencies of neutrophils (NE), monocytes (Mono), and lymphocytes (Lymph) in the peripheral blood (PB) of WT and *Pstpip2^−/−^* mice (n≥5 per group). (**H**) Representative IHC images of Ly6G on hind paw sections from WT and *Pstpip2^−/−^* mice. Stain in brown: neutrophil marker Ly6G; scale bar, 200 μm. (**I**) Principal component analysis (PCA) of WT and *Pstpip2^−/−^* mice by bulk RNA-seq. (**J**) Volcano plots of DEGs in BM cells of WT and *Pstpip2^−/−^* mice by bulk RNA-seq. (**K**) Representative micro-CT of hind paw bones from 20-week-old WT and *Pstpip2^−/−^* mice. Gray images show total bone tissue while pseudo-color images provide difference between high-density (in yellow) and low-density (in blue) area. (**L**) Measurement and quantification of superoxide in WT and *Pstpip2^−/−^* mice. The mice were injected (i.p.) with L-012 chemiluminescent probe before imaging. Two-tailed Student’s t test, mean±SEM; \**p*<0.05, \*\*\**p*<0.001, \*\*\*\**p*<0.0001.

To investigate the alteration of immune cells in *Pstpip2^−/−^* osteomyelitis, the number and frequency of immune cells were characterized. The hind paws of WT and *Pstpip2^−/−^* were isolated and examined through hematoxylin and eosin (H&E) staining. *Pstpip2^−/−^* mice showed very high levels of immune cells and granulocyte infiltration in the hind paws, whereas WT mice showed no such infiltration (Fig. 1F). By collecting peripheral blood samples from the two groups of mice for hemogram detection, we found that the frequency of neutrophils and monocytes in the peripheral blood of *Pstpip2^−/−^*mice was significantly increased, while the frequency of lymphocytes was inversely decreased (Fig. 1G). The results of immunohistochemical (IHC) staining showed that *Pstpip2^−/−^* mice had obviously abundant neutrophils in the hind paw tissue compared to WT (Fig. 1H). These data suggest that myeloid cells represented by neutrophils and monocytes were likely the major cell types that drive osteomyelitis caused by *Pstpip2* mutation or deficiency.

Next, we performed bulk RNA sequencing (RNA-seq) on BM cells from WT and *Pstpip2^−/−^*mice to compare the differences at the transcriptome level (Fig. 1I). Analysis of differential expressed genes (DEGs) showed that *Pstpip2^−/−^* mice had significantly increased mRNA expression levels of cytokines compared to WT, including *Il1b* and *Ccl3* (Fig. 1J). There were studies showing that cytokines such as IL-1β and CCL3 (MIP-1α) acted as osteoclasts-activating factors and promoted the dissolution of bone matrix in a variety of diseases^[25–30]^. The classic inflammatory phenotype of CRMO is accompanied by severe bone damage. Therefore, we focused on the differences in bone tissue structure between the two groups of mice. Microscopic CT (micro-CT) results showed that compared with WT mice, there were multiple low-density areas in the hind paw bone tissue of *Pstpip2^−/−^* mice, suggesting severe damage to the bone tissue (Fig. 1K). Phagocytes (especially neutrophils) are the central of inflammatory response, and nicotinamide adenine dinucleotide phosphate oxidase (NADPH oxidase) present in them is one of the major sources of reactive oxygen species (ROS)^[31, 32]^. ROS not only involve in regulating cell growth, differentiation, apoptosis, and activation of cell signaling cascades as intracellular second messengers, but also mediate inflammation by oxidizing protein and lipid cellular components and damaging DNA^[32, 33]^. Moreover, ROS are also important components in regulating OC differentiation and activation^[33]^. We therefore visualize ROS production in living mice by using luminol derivative L-012. We observed a more intense luminescence signal in the hind paws of *Pstpip2^−/−^* mice compared with that of WT mice (Fig. 1L), suggesting that ROS overproduction may be partially involved in the genesis of inflammation and osteolysis. Taking these results together, we demonstrated that the 5-bp deletion in *Pstpip2* induces a CRMO-like phenotypes, characterized by progressive swelling of the hind paws accompanied by severe bone damage. The above results suggest that the occurrence and aggravation of this CRMO-like disease are strongly correlated with the increase in the frequency of myeloid cells, the level of inflammatory cytokines and the production of ROS.

### Deletion of *Morrbid* significantly inhibits the CRMO-like disease progression caused by the 5-bp deletion in *Pstpip2*

*Morrbid* is a long non-coding RNA (lncRNA) that is specifically highly expressed in mature, short-lived myeloid cells and is conserved between humans and mice. We and others have shown that *Morrbid*-deficient mice exhibit a drastic reduction in mature myeloid cells, such as eosinophils, neutrophils, and monocytes, in blood and tissues. By regulating its downstream pro-apoptotic gene *Bcl2l11* in *cis*, *Morrbid* regulates apoptosis and controls the lifespan of these cells^[19]^. We therefore asked about the expression levels of both *Morrbid* and *Pstpip2* in murine myeloid cells. Using scRNA-seq, we observed that both *Morrbid* and *Pstpip2* were specifically highly expressed in myeloid cells such as Mono, GMP, and NE (Fig. 2A).

**Figure 2.**
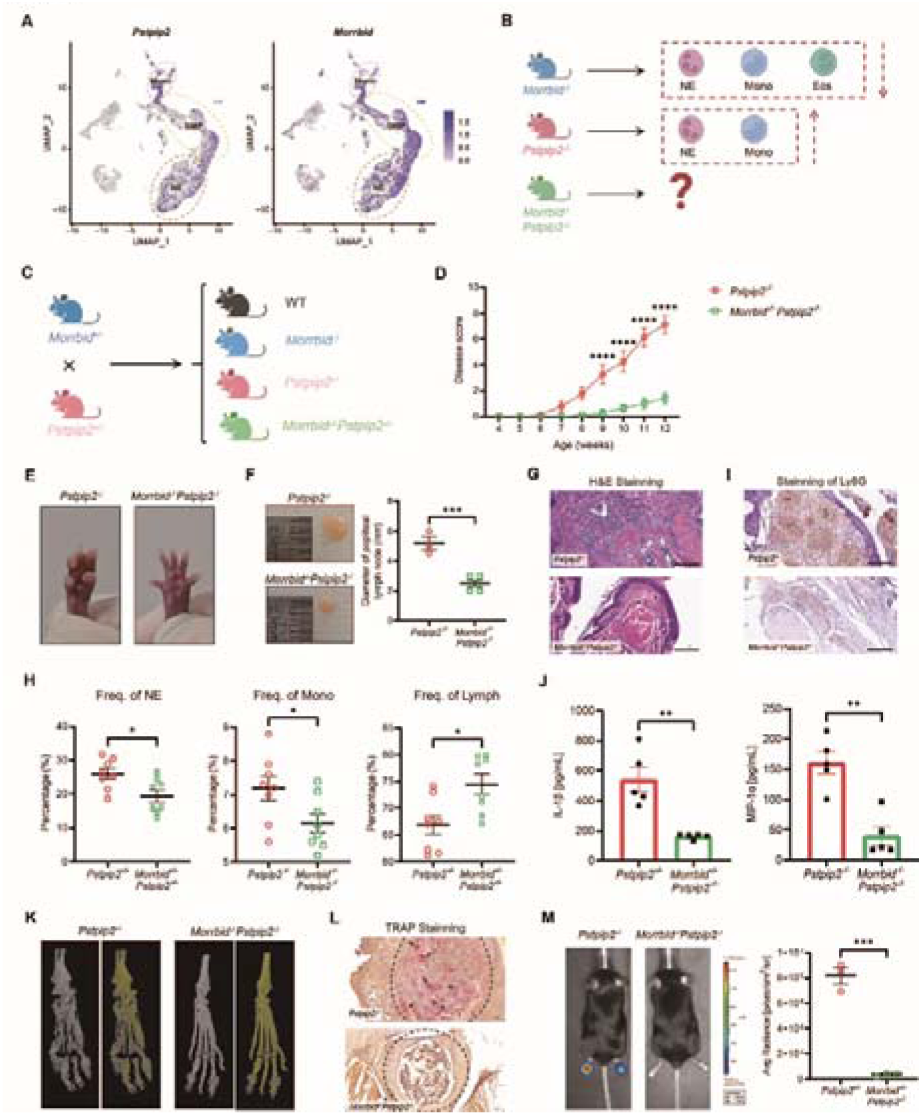
Loss of *Morrbid* significantly inhibits the CMO disease progression. (**A**) Expression of *Pstpip2* and *Morrbid* in the scRNA-seq dataset of BM cells from hind paws in WT mice. Mono, monocytes; Neu, neutrophils; GMP, granulocyte-macrophage-progenitors. (**B**) Schematic for the rationale and hypothesis in this study. Red dashed boxes indicate the relevant major myeloid cell types, and red dashed arrows indicate increased or decreased cell number and/or proportion in mutants compared to the WT mice. (**C**) Schematic for the construction of the compound mutant *Morrbid^−/−^Pstpip2^−/−^* mice. (**D**) Disease severity was scored (scale from 0 to 10) by visual inspection of the hind paws of *Pstpip2^−/−^* and *Morrbid^−/−^Pstpip2^−/−^*mice. n=8 per group. (**E**) Representative images of the hind paws of 20-week-old *Pstpip2^−/−^* and *Morrbid^−/−^Pstpip2^−/−^* mice. (**F**) Representative images and quantification (size in diameter) of the popliteal lymph nodes of 20-week-old *Pstpip2^−/−^* and *Morrbid^−/−^Pstpip2^−/−^* mice. n=3 for *Pstpip2^−/−^*, n=4 for *Morrbid^−/−^Pstpip2^−/−^*. (**G**) Representative H&E images of hind paw sections of *Pstpip2^−/−^* and *Morrbid^−/−^Pstpip2^−/−^* mice. scale bar, 200 μm. (**H**) Frequencies of Neu, Mono, and Lymph in the peripheral blood of *Pstpip2^−/−^* and *Morrbid^−/−^Pstpip2^−/−^* mice. n=8 per group. (**I**) Representative IHC images of hind paw sections of *Pstpip2^−/−^* and *Morrbid^−/−^Pstpip2^−/−^* mice. Stain in brown: neutrophil marker Ly6G; scale bar, 200 μm. (**J**) IL-1β and MIP-1α (CCL3) concentration in hind paw lysates detected by ELISA. n=5 per group. (**K**) Representative micro-CT of hind paw bones from 20-week-old *Pstpip2^−/−^* and *Morrbid^−/−^Pstpip2^−/−^* mice. Gray images show total bone tissue while pseud-ocolor images provides difference between high-density (in yellow) and low-density (in blue) area. (**L**) Representative histologic sections of hind paws from *Pstpip2^−/−^* and *Morrbid^−/−^Pstpip2^−/−^*mice stained by TRAP (a marker for osteoclasts; arrows), the dashed line marks BM cavity region. scale bar, 100 μm. (**M**) Measurement and quantification of superoxide in *Pstpip2^−/−^* and *Morrbid^−/−^Pstpip2^−/−^*mice. The mice were injected (i.p.) with L-012 chemiluminescent probe before imaging. Two-tailed Student’s *t*-test, mean±SEM;\**p*<0.05,\*\**p*< 0.01,\*\*\**p*<0.001,\*\*\*\**p*<0.0001.

In view of the massive expansion of myeloid cells in the peripheral blood and hind paws sites of the *Pstpip2^−/−^* mice, we hypothesize that *Morrbid* deletion retards progression of CRMO-like syptoms (Fig. 2B). We investigated whether ablation of *Morrbid* in the *Pstpip2^−/−^* mice would be sufficient to observe the rescue, and thus, DKO mice were generated by crossing *Morrbid^+/−^* mice with *Pstpip2^+/−^* mice. *Pstpip2^−/−^* mice were used as controls for further study of the function of *Morrbid* in osteomyelitis (Fig. 2C). We found that *Morrbid* knockout significantly attenuated and delayed the development of hind paw swelling caused by *Pstpip2* depletion using our scoring system (Fig. 2D). Furthermore, the effect was durable rather than temporary, as evidenced by our observations in the hind paws of two groups of 20-week-old mice (Fig. 2E). We dissected popLN of both groups of mice, measured diameters and performed statistics, and found that *Morrbid^−/−^Pstpip2^−/−^* mice had apparently smaller popliteal lymph nodes compared with *Pstpip2^−/−^* mice, suggesting that inflammation in the surrounding tissues was controlled (Fig. 2F). These phenotypes indicated that deletion of *Morrbid* effectively rescued the development of inflammatory response in CMO mice.

We next addressed the effects of *Morrbid* on immune cells in CMO. H&E staining of the mice hind paws showed that *Morrbid^−/−^Pstpip2^−/−^* mice had significantly reduced infiltration of immune cells and inflammatory granulomas (Fig. 2G). The hemogram analysis for peripheral blood from the two groups of mice showed that the *Morrbid^−/−^Pstpip2^−/−^* mice had a significantly reduced frequency of neutrophils and monocytes and increased frequency of lymphocytes as compared with the *Pstpip2^−/−^* mice (Fig. 2H). IHC staining of the hind paws revealed a significant reduction in the number of neutrophils in *Morrbid^−/−^Pstpip2^−/−^* mice (Fig. 2I). The excessive activation of immune cells releases a mass of cytokines and enhances the self-feedforward of inflammation^[34]^. The hind paw homogenates of the two groups of mice were obtained for enzyme linked immunosorbent assay (ELISA) with the aim of measurement of the protein levels of IL-1β and MIP-1α (CCL3) in hind paws. The results showed that the two cytokines from *Morrbid^−/−^Pstpip2^−/−^* mice were all significantly lower than that from *Pstpip2^−/−^* mice (Fig. 2J). These data suggest that effective reduction of inflammation in the hind paws of CRMO-like mice could be achieved by *Morrbid* deletion, and the reduction of myeloid cells and cytokines in the lesions are important for mitigating the CRMO-like symptoms.

In addition to this, we also quantified bone damage in the DKO mice. By micro-CT, we confirmed that knockout of *Morrbid* in *Pstpip2^−/−^* mice resulted in an evident reduction in low-density areas and significantly decreased the degree of osteolysis (Fig. 2K). Bone tissue homeostasis is synergistically maintained as supported by osteoblastic bone formation and osteoclastic bone resorption^[35]^. Tartrate-resistant acid phosphatase (TRAP), as a specific marker of osteoclasts, plays a critical role in the synthesis and degradation of bone tissue^[36–39]^. By staining TRAP in hind paw sections from both groups of mice, we found that *Morrbid^−/−^Pstpip2^−/−^* mice had a decreased number of osteoclasts and an increased complete bone marrow cavity in the hind paw accordingly (Fig. 2L). These results suggest that the rescue of bone damage in *Pstpip2^−/−^* by *Morrbid* deletion is likely due to inhibition of OC formation. We also evaluated the effect of *Morrbid* deletion on ROS production. We observed a significant reduction in ROS luminescence signal in *Morrbid^−/−^Pstpip2^−/−^* mice compared with that in *Pstpip2^−/−^* mice after intraperitoneal injection of L-012 (Fig. 2M). In conclusion, deletion of *Morrbid* significantly reduced the frequencies of myeloid cells and the levels of inflammatory cytokines in the hind paws of CMO mice, which effectively inhibited the symptoms and progression of osteomyelitis.

### sc-RNA seq reveals the pathogenesis of *Pstpip2*-deficient CRMO and that loss of *Morrbid* relieves autoinflammatory symptoms in *Pstpip2^−/−^* mice

scRNA-seq analysis was performed on hind paws of four groups of mice (WT, *Morrbid^−/−^*, *Pstpip2^−/−^* and DKO). Muscle cells were removed and only cells from the bone marrow were included for the library construction and sequencing procedure (see methods for details). After standard Seurat annotation pipeline and filtering mesenchymal cells, around 8,000 to 10,000 hematopoietic cells (CD45^+^) per group were included for further annotation analysis after quality control. Annotation of cells by canonical cell marker genes grouped all cells into 11 cell populations, including GMP, common lymphoid precursors (CLP), hematopoietic stem and progenitor cells (HSPC), and Plasmablasts, monocytes, neutrophils, osteoclasts, mast cells, plasmacytoid dendritic cells (pDC), and lymphoid cells (T and B cells) (Fig. 3A, B, Supplemental Figure 1A).

**Figure 3.**
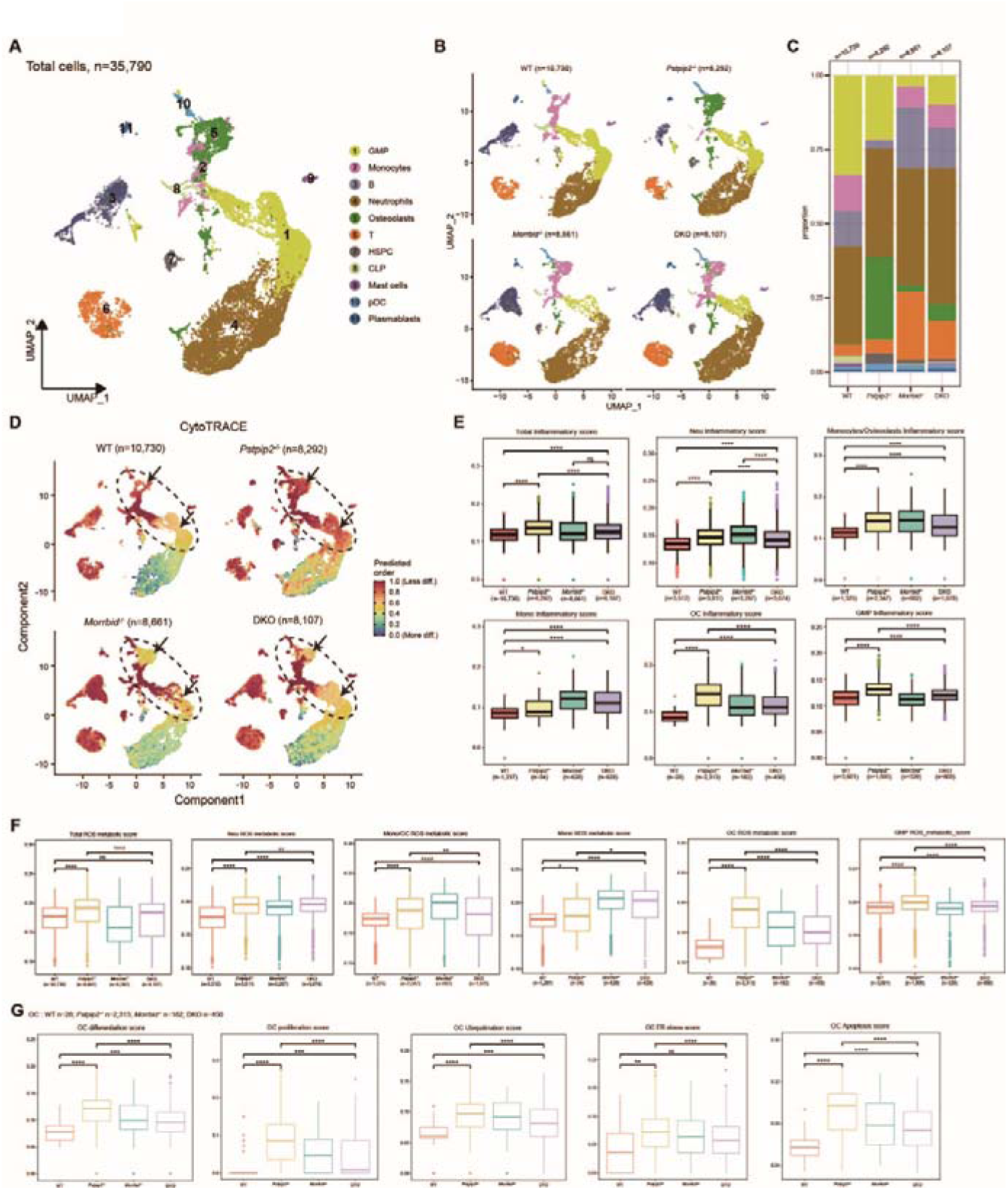
scRNA-seq analysis of the BM cells from the hind paws of four genotype mice. (**A**) Uniform manifold approximation and projection (UMAP) visualization of the BM cells (n=35,790); each dot represents a cell, and a color indicates a cell-type as indicated. **(B**) Separate UMAP plots of BM cells from the four genotype mice. (**C**) Stacked chart of the proportion of each cell-type in the four genotype mice. (**D**) Heatmap for the stemness scores of BM cells from the four groups of mice by CytoTRACE. Higher predicted values (dark red) suggest state with higher stemness (less differentiated and more progenitor-like). Note that monocytes, neutrophils and GMP from *Pstpip2^−/−^* appear more immature compared to the myeloid cells form other three genotypes. (**E** and **F**) The total and various types of myeloid cells inflammatory gene signature scores and ROS metabolic gene signature scores of the BM cells from four groups of mice. Myeloid-lineage cells including GMP, monocytes, neutrophils, osteoclasts were included for scoring the total score. The individual scores were also computed as indicated. (**G**) Utilizing the scRNA-seq datasets we also measured the activity of osteoclasts by computing differentiation score, proliferation score, ubiquitination score, endoplasmic reticulum stress score and apoptosis score of osteoclasts from the four groups of mice.

Datasets from each genotype group had all of these cell populations, but the overall proportion of myeloid cells was higher in *Pstpip2^−/−^* mice than in WT and in *Morrbid^−/−^ Pstpip2^−/−^* mice (Figure 3C). We asked the effect of *Pstpip2* mutation and *Morrbid* deletion on the development states of different cell types. We visualized the cellular stemness of individual cells by using cellular trajectory reconstruction analysis (CytoTRACE) using gene counts and expression. Results show that the OC, GMP and Neu prediction score in *Pstpip2^−/−^* mice are higher than that in WT and DKO mice, suggesting their stemness level are enhanced (Figure 3D). These cells from *Pstpip2^−/−^*were at a more primitive level, suggesting the time intervals before the occurrence of apoptosis were likely larger, and their survival time may be extended. However, loss of *Morrbid* predisposed these myeloid cells to undergo apoptosis earlier, as supported by the lower stemness scores of these cells in DKO mice (Figure 3D).

Consistent with results in Fig. 1l and Fig. 2m, *Pstpip2^−/−^* mice has higher inflammatory and ROS metabolic scores on total cells and all myeloid cell types than WT and DKO mice (Figure 3, E-F). Compared with WT and DKO, *Pstpip2 ^−/−^* has higher differentiation score, proliferation score, ubiquitination score, endoplasmic reticulum (ER) stress score and apoptosis score (Figure 3G). We performed analyses of DEGs in cell clusters between different groups. The results showed that matrix metalloproteinase genes (*Mmp3* and *Mmp13*) and serum amyloid gene (*Saa3*) were upregulated in multiple cell populations from *Pstpip2^−/−^* mice compared to that from WT mice (Supplemental Figure 1B). And high expression of these genes was positively correlated with inflammatory disease progression^[40–45]^. In contrast, the expression of these genes was significantly reduced in DKO mice (Supplemental Figure 1C). However, lncRNA Malat1 and the proteoglycan 4 gene (*Prg4*) were up-regulated in DKO mice (Supplemental Figure 1C). Multiple studies have shown the role of lncRNA Malat1 in bone and cartilage diseases and osteogenic differentiation^[46–48]^. Proteoglycan encoded by *Prg4* was reported to function as a cartilage surface lubricant and involve in repairing articular cartilage damage in mouse models of inducible full-thickness injury^[45, 49]^. These results suggest that the expression of these genes is upregulated in *Pstpip2^−/−^* mice, and loss of *Morrbid* inhibits the expression level of these genes and alleviates the symptoms.

We then focused on three cell types from the scRNA-seq datasets, GMP, OC, and NE. These cells were specifically extracted and clustered again to distinguish their subsets. Based on the expression of typical marker genes, we annotated a total of four cell subsets in GMP, four cell subsets in OC, and five cell subsets in NE (Figure 4A①, 4B ①, 4C①). Gene Ontology (GO) enrichment analysis was used for all cell subsets to understand the function of each cell type. GMP seemed to be related to the catabolism of collagen. OC was highly associated with the response to IL-1 and reactive oxygen species metabolism process. The chemotaxis-related functions of NE were significantly enriched (Figure 4A②, 4B②, 4C②). These biological functions suggest a key role for these three cell types in CRMO-like disease progression.

**Figure 4.**
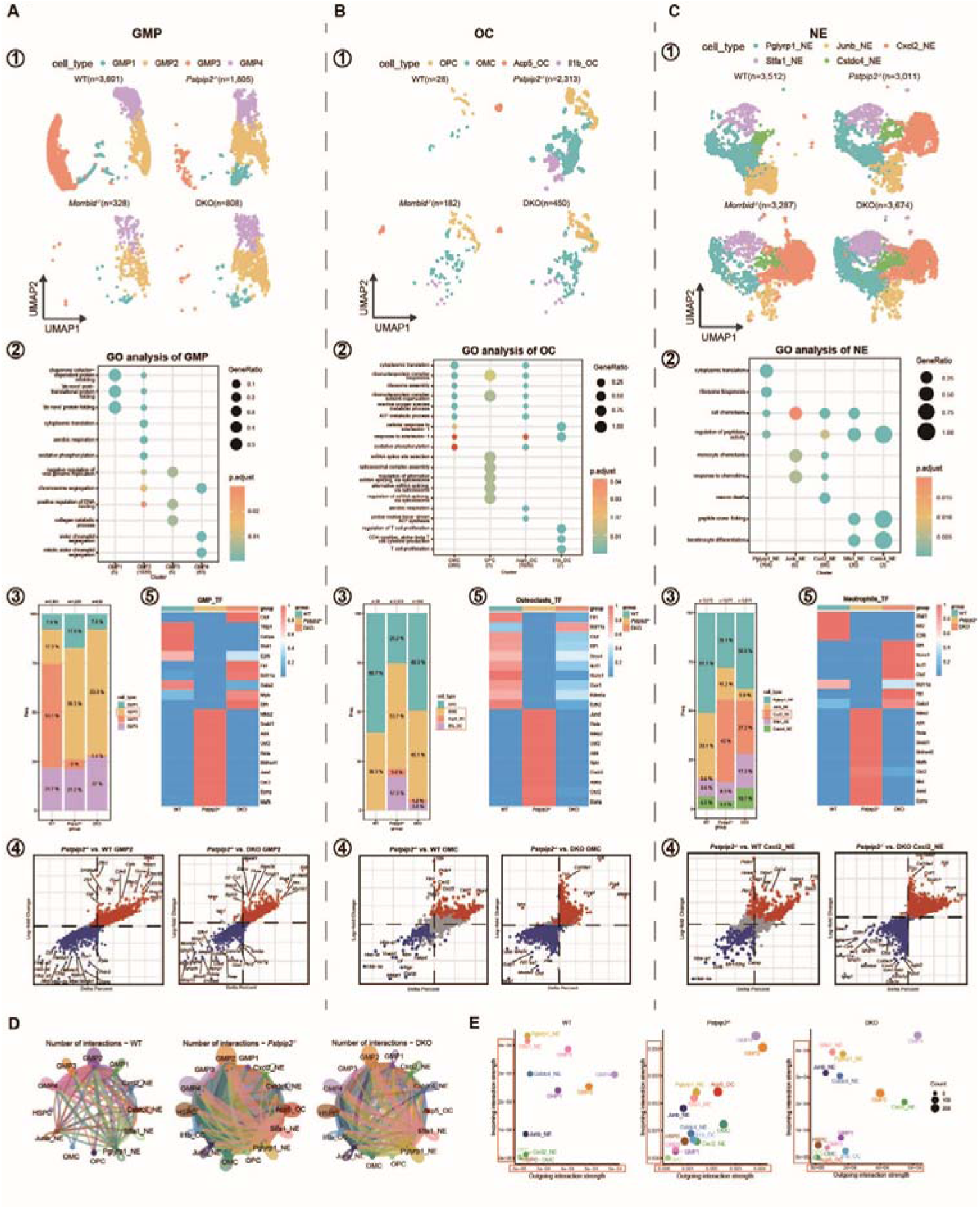
Characteristics of inflammatory cells and osteoclasts in *Pstpip2^−/−^* and *Morrbid^−/−^Pstpip2^−/−^* mice. (**A-C**) Myeloid-linage cells including GMP, OC and NE were extracted for further analysis. Four genotype mice were included: WT, *Pstpip2^−/−^*, *Morrbid^−/−^* and DKO. GMP, OC and NE cells from Figure 3A were extracted for further analysis as shown. ①UMAP plots of GMP, OC and NE respectively;②GO enrichment analysis of different cell types as indicated; ③Stacked charts of the various cell subsets proportion in GMP, OC and NE respectively;④Scatter plots showing DEGs in the indicated comparison. Each point represents a gene;⑤Heatmap of TF-RAS in GMP, OC and NE respectively. (**D**) Circle plots showing cell-cell communications between cell subtypes in the BM scRNA-seq datasets from WT, *Pstpip2^−/−^*, and DKO mice. (**E**) Overall signal intensities based on Outgoing (x-axis) and Incoming (y-axis) strengths. Of note, a much higher basal level of signals was determined in *Pstpip2^−/−^*(∼10-fold higher).

According to the comparison of the proportion of each cell subtype between WT and *Pstpip2^−/−^* mice, we paid attention on the cell subtypes with large differences between these two groups, namely GMP2, OMC and Cxcl2_NE (Figure 4A③, 4B③, 4C③), and carried out further analysis. The results of differential gene analysis between different groups of these cell subsets showed that inflammation and peroxidation related genes, such as *Saa3*, *Grina*, *Mmp13* and *Prdx1*, were significantly upregulated in *Pstpip2^−/−^*. Oppositely, the expression of genes involved in bone protection and remodeling, such as *Suco* and *Prg4*, were significantly higher in WT and DKO (Figure 4A④, 4B④, 4C④). Kyoto encyclopedia of genes and genomes (KEGG) enrichment analysis of GMP2 and Cxcl2_NE and GO analysis of OMC showed that the pathways and functions of cell metabolism, rheumatoid arthritis and osteoclast differentiation were more significant in *Pstpip2^−/−^* compared with WT and DKO (Supplemental Figure 1, D-E). These results further supported a mechanism by which loss of *Pstpip2* induces CMO and loss of *Morrbid* inhibits CRMO-like progression. The *Pstpip2^−/−^* had Il1b_OC and Acp5_OC, which were not present in the WT, and the proportion and number of these two OC subsets in DKO were obviously lower than those in *Pstpip2^−/−^*. We therefore inferred that these two subsets of OC may be the major osteoclast population that play an osteolytic role in CMO (Figure 4A③, 4B③, 4C③).

The differences between the two subtypes of OC were analyzed in the comparison between *Pstpip2^−/−^* and DKO. However, Acp5_OC in the DKO could not be enriched due to the few numbers of cells. Then we compared the differences in Il1b_OC. DEG analysis showed that inflammation-related genes, such as *Mmp13*, *Grina* and *S100a8*, were highly expressed in *Pstpip2^−/−^*, and osteoprotective genes, *Prg4*, were significantly upregulated in DKO (Supplemental Figure 1F). GO enrichment analysis showed that OC in this subtype was associated with immune disorders and the emergence of some chronic inflammatory diseases (Supplemental Figure 1F). All the above results suggest that the proportion and function changes of myeloid cells play prominent roles in the progression of CRMO-like symptoms in *Pstpip2^−/−^*, and DKO mice reverses these effects.

We next discuss the states of cell-to-cell interactions between different groups in these cell types using CellChat. The number of interactions was greater in the *Pstpip2^−/−^* than in the other two groups (Figure 4D). Comparison of the number and strength of intercellular interactions between *Pstpip2^−/−^* and DKO mice (DKO vs. *Pstpip2^−/−^*) showed that most intercellular interactions were enhanced in *Pstpip2^−/−^* (in blue). Only a few of interactions were enhanced in DKO (in red) (Supplemental Figure 1G). These results suggest that cell activity is higher in CRMO-like diseased tissues. In the pairwise comparison of the overall signaling patterns, the signaling pathways with differential intensities were mainly affected in Acp5_OC, including MIF, VISFATIN, SPP1, CSF3, TGFβ and GDF signaling pathways. These signaling pathways were significantly enhanced in Acp5_OC of *Pstpip2^−/−^* (Supplemental Figure 1H). Comparing the differences in GMP signaling pathways among the three groups, we found that TGFβ, SEMA3, KIT, APRIL and BAFF signaling pathways were more intense in DKO (Supplemental Figure 1I). In addition, the intensities of the incoming and outgoing interactions as a whole in *Pstpip2^−/−^* were higher than that in WT and DKO. The interaction strength of NE of the various cell types in the DKO was similar to that in WT, and the interaction strength of OC of the various cell types in the DKO was lower than that in *Pstpip2^−/−^* (Figure 4E). These results show that intercellular communication is enhanced in CMO, especially in some signaling pathways related to myeloid cell activation and inflammation occurrence.

By visualizing the transcriptional factor regulon activity score (TF-RAS) of different cell types in each group, we explored the transcription factor activities of different cell types in three groups. We found that some transcription factors that are usually highly expressed in inflammatory diseases, Nfkb2, Rela, Jund, Atf4 and Esrra, had higher scores among the three cell types of *Pstpip2^−/−^*; Ctcf, Elf1, Fli1, Bcl11a, and Runx1 had higher TF-RAS in the WT and DKO (Figure 4A⑤, 4B⑤, 4C⑤), which are partially associated with bone protection in some diseases. Similar results were observed for the distribution and expression levels of these transcription factors (Supplemental Figure 1J). These results suggest that the pathogenesis of CRMO-like autoinflammation may be associated with the changes of some transcription factors and the activation of some pathways (e.g., NF-κB).

Taken together, the inter-cellular mechanism of CRMO caused by *Pstpip2* mutation is complicated, involving various aspects such as the level of gene expression, transcription factors and developmental stage regulation. Loss of *Morrbid* regulates myeloid cells from multiple points, and ultimately inhibits inflammation progression and bone damage in CMO.

### Experimental validation in the four genotype mice

Based on the results of the above, we have an unbiased and comprehensive understanding of pathology of CRMO-like diseases and the involvement of *Morrbid* in the disease. To verify our results concluded above, we further performed experiments including the flow cytometry and qRT-PCR. We extracted whole bone marrow cells from the four genotype mice (WT, *Morrbid^−/−^*, *Pstpip2^−/−^* and DKO) and measured the frequencies of different types of myeloid cells and osteoclast precursors (OCP) in bone marrow by flow cytometry. Compared with WT and *Morrbid^−/−^Pstpip2^−/−^* mice, the frequencies of myeloid cells (Gr1^+^CD11b^+^), monocytes (c-Kit^+^/c-Kit^−^CD11b^+^Ly6C^+^), and OCP (c-Kit^+^CD11b^lo^Csf1r^+^) were all significantly higher or tended to be higher in the bone marrow of *Pstpip2^−/−^* mice (Figure 5A-D). A similar trend was observed in neutrophils (c-Kit^+^/c-Kit^−^CD11b^+^Ly6G^+^) (Figure 5B-C). These results are consistent with the previous results showed in scRNA-seq dataset that elevated myeloid cell frequencies may contribute to CRMO-like diseases and that loss of *Morrbid* inhibits CRMO progression by repressing such elevations.

**Figure 5.**
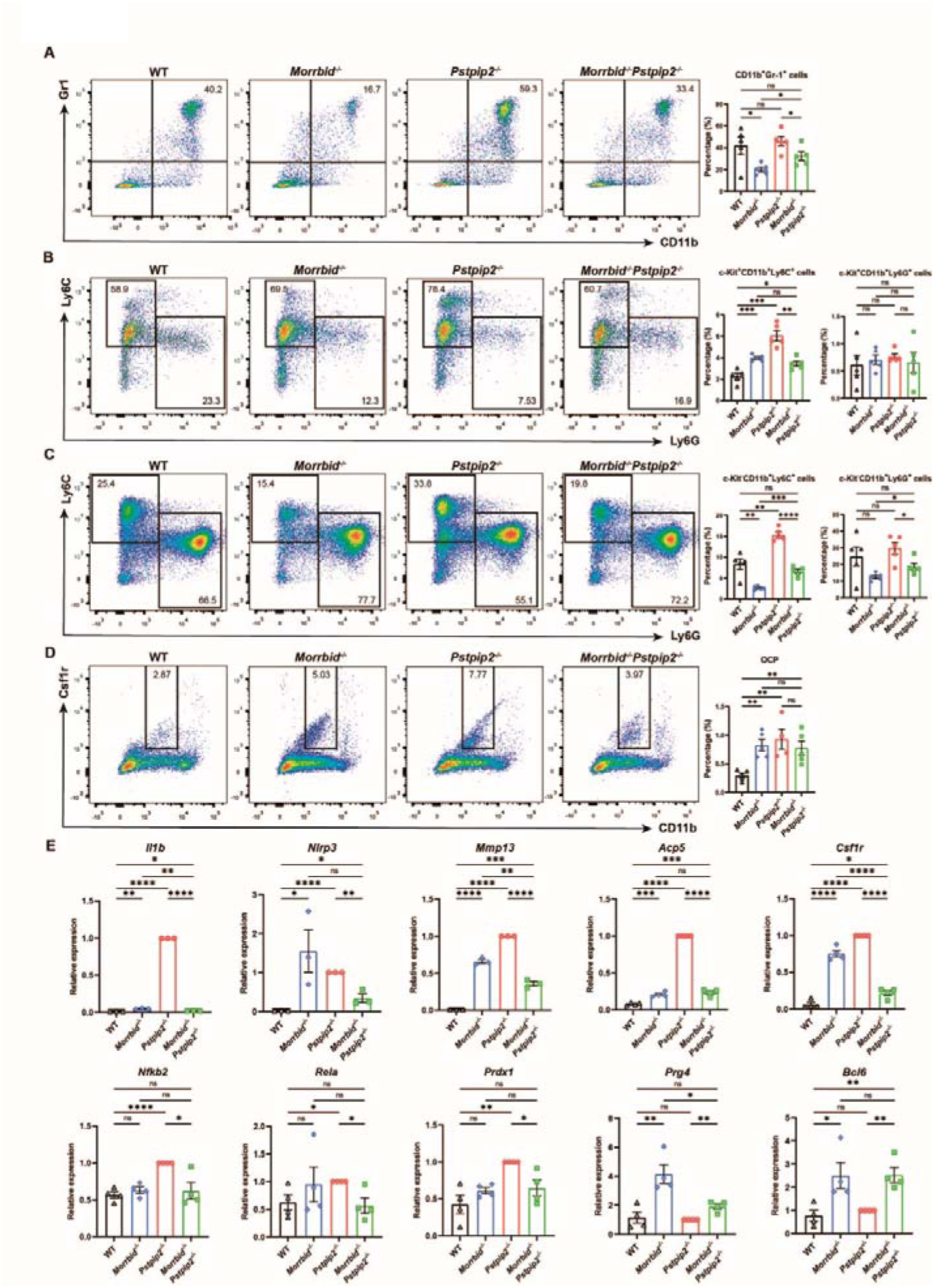
Experimental analysis using flow cytometry and qRT-PCR for validating findings from scRNA-seq analysis. (**A-D**) Representative flow cytometry profiles in BM samples from the four groups of mice (left) and quantification of the results (right). (**A**) myeloid cells (Gr1^+^CD11b^+^); (**B**) pre-mature monocytes (c-Kit^+^CD11b^+^Ly6C^+^) and neutrophils (c-Kit^+^CD11b^+^Ly6G^+^); (**C**) mature monocytes (c-Kit^−^CD11b^+^Ly6C^+^) and neutrophils (c-Kit^−^CD11b^+^Ly6G^+^); (**D**) osteoclast precursors (c-Kit^+^CD11b^low^Csf1r^+^). n≥4 animals for each group. (**E**) The expression levels of indicated genes in hind paw lysates, determined by qRT-PCR. n≥3 animals for each group. Two-tailed Student’s t test, mean ± SEM; \**p* ≤ 0.05, \*\**p* ≤ 0.01, \*\*\**p* ≤ 0.001, \*\*\*\**p* ≤ 0.0001, ns, not statistically significant.

We extracted total RNA from the hind paws of mice with different genotypes and measured expression levels of some genes by qRT-PCR. The level of *Il1b* expression in the hind paw of mice was similar to our previous data for IL-1β (Figure 5E). There was a study shown that NLRP3 inflammasome is involved in the pathogenesis of CRMO-like disease in *Pstpip2^cmo/cmo^*, not only by converting pro-IL-1β to active IL-1β, but also by ROS production and cellular stress^[14]^. In our data analysis of cellular transcription factors, we also found that transcription factor Nfkb2, which is related to the NLRP3 pathway, was significantly active in the tissues of *Pstpip2^−/−^*mice hind paws. The expression level of *Nlrp3* in our four groups of mice was measured and confirmed that its important role in CRMO (Figure 5E). We also examined the expression levels of other inflammation-related genes (e.g., *Mmp13*, *Acp5*, *Csf1r*, *Prdx1*, *Prg4*) and gene encoding representative transcription factors (e.g., *Nfkb2*, *Rela*, *Bcl6*) identified by scRNA-seq. These results were similar to those obtained with single-cell transcriptome analysis, which indicate that loss of *Morrbid* alleviated the inflammatory symptoms and bone tissue damage in CRMO. This alleviation is associated with down-regulation of some genes within the NF-κB signaling pathway that accelerates inflammatory progression.

## Discussion

Chronic non-bacterial osteomyelitis is an inflammation of bone tissue without bacterial infection. When it is active and occurs in multiple foci, it is referred to as CRMO^[50]^. Patients with CRMO usually present with recurrent fever and bone pain, and swelling and deformity may occur in the affected area^[8]^. Here we described a new *Pstpip2* mutant mouse strain with 5-bp loss in the protein-coding region, interrogated the pathogenesis of CRMO-like phenotypes by scRNA-seq analysis, and demonstrated the efficacy of targeting *Morrbid* for mitigating CRMO. In summary, our findings provided in-depth insights into the relevance and function of lncRNA *Morrbid* in the context of AID and CRMO, and demonstrated *Morrbid* is a target for inhibiting osteomyelitis.

PSTPIP2, an adaptor protein that located in myeloid cell membranes, regulates membrane curvature and effects the morphology of cell membrane, stabilizing the cytoskeleton and regulating the biological functions of myeloid cells, including endocytosis, podosome formation, cell division and movement. PSTPIP2 negatively regulates the activation of phagocytes and inhibits the occurrence of inflammation. In osteoclasts, PSTPIP2 promotes the decomposition of osteoclasts and regulates the dynamic homeostasis of osteoblasts and osteoclasts^[15]^. In previous studies, the *cmo* mutation in *Pstpip2* inactivates the PSTPIP2 protein, resulting in uncontrolled bone-destroy functions of OC. Mice with *Pstpip2* mutation develop CRMO lesions and show a massive increase in myeloid cells at the lesion location. The overproduction of osteoclasts causes bone damage at the lesion site and severely destroys the basic structure of bone. Massive infiltration of neutrophils and monocytes/macrophages leads to excess release of proinflammatory cytokines. In addition, the abundant NADPH oxidase in phagocytes produces a large amount of ROS, which further promotes the progression of inflammation and osteolysis. Biased myelogenesis is also indicated the perpetual cycle for the progression of CRMO in the *Pstpip2^cmo/cmo^*. However, the efficacy of depleting inflammatory cells in CRMO has not been tested until this study.

Non-steroidal anti-inflammatory drugs (NSAIDs) are commonly used as early treatment for CRMO^[11, 51]^. When inflammation continues to worsen, off-label empirical therapies, including bisphosphonates and TNF-α antagonists, are used^[11, 51]^. However, there are significant differences in the response to these drugs among different patients^[51]^. There is currently no standardized treatment regimen and no FDA-approved drug for the treatment of CRMO^[8, 11]^. Therefore, it is imperative to develop effective CRMO inhibition methods. *Morrbid* as a lncRNA controls myeloid cell lifespan. Myeloid cells deficient in *Morrbid* exhibited increased apoptosis. Our experiments confirmed that deletion of *Morrbid* in *Pstpip2^−/−^* mice significantly reduced the number of myeloid cells at the lesion site, lowered inflammatory cytokines and ROS levels, and alleviated bone damage. The reduced expression of *Morrbid* suggests control over the pathogenic mechanisms of CRMO, ultimately suppressing inflammation and bone damage symptoms associated with CRMO disease. The high conservation of *Morrbid* between humans and mice suggests the possibility of *Morrbid* as a target to inhibit the development of the disease. How to effectively and specifically target *Morrbid* will be a challenge to be overcome. In addition, the main cell type driving the initiation of CRMO in *Pstpip2^−/−^* mice (or *Pstpip2^cmo/cmo^*) remains controversial. The prominent role of myeloid cells in the progression of CRMO has been observed in previous studies and this study. Previous studies held the view that neutrophil or monocyte lineage drives the initiation of CRMO^[11, 14, 17, 52]^. Our study together with single-cell RNA sequencing data confirmed an increased frequency of multiple types of myeloid cells, especially neutrophils and osteoclasts, in the lesions and peripheral blood of mice with CRMO.

By scRNA-seq of the four genotype mice, *Pstpip2^−/−^* mice showed increased inflammatory levels and ROS production, significantly upregulating the expression of genes that promote inflammation. Furthermore, *Pstpip2^−/−^* mice enhanced intercellular communication and increased the activity of transcription factors related to NK-κB pathway. The deficiency of *Morrbid* resulted in a comprehensive inhibitory effect in these aspects. However, further studies are required to determine which type of myeloid cell plays a dominant role in the CRMO initiation, and the primary cell type exerts inhibitory effects when deletion of *Morrbid*. Based on the experimental results and single-cell RNA-seq data analyses, we tend to suggest that neutrophils play a key role in the process of the inflammation aggravation. The excess release of inflammatory cytokines further promotes the activation of monocytes into osteoclasts, leading to osteolysis.

PSTPIP2 binds to its interaction partners and plays a key role in membrane ligation processes in eukaryotic cells^[15]^. It is widely involved in cell endocytosis, division and morphological regulation^[15, 16, 53]^. It is mainly expressed in cells of the monocyte or macrophage lineage, and is highly relevant to the migration of macrophages and the decomposition of osteoclasts^[15]^. *Morrbid* not only has the ability to regulate myeloid cell apoptosis^[19]^, but also has an expression profile highly similar to *Pstpip2* in mice. Importantly, the study of *Morrbid* in osteoclast formation and function is still lacking now. Our study fills a gap in this area. However, the mechanism by which *Morrbid* functions and its effects on PSTPIP2 interaction partners need to be further explored.

## Methods

### Animals

We used wild-type (WT), *Morrbid^−/−^*, *Pstpip2^−/−^*, and *Morrbid^−/−^Pstpip2^−/−^* mice in this study. All mice were on C57BL/6 background. *Morrbid^−/−^* and *Pstpip2^−/−^* mice were generated by and purchased from Cyagen Biosciences (Suzhou, China). *Morrbid^−/−^* mice were generated by knockout of six exons of *Morrbid*. *Pstpip2^−/−^* mice were generated by a 5-bp deletion in *Pstpip2* exon 2. *Morrbid^−/−^Pstpip2^−/−^*mice were generated by crossing *Morrbid^−/−^* mice with *Pstpip2^−/−^* mice. All animal experimentations were performed in accordance with protocols approved by the Animal Care and Use Committee of Tianjin Medical University.

### Histopathology

The hind paws were fixed in 4% paraformaldehyde (#p1110, Solarbio) for 24 h and decalcified in EDTA Decalcified Solution (#BL616B, biosharp) for 2 to 3 weeks, followed by paraffin embedding and histological cutting. The slice thickness was 4 μm. Hematoxylin and eosin (H&E) staining was used to detect the infiltration of immune cells and inflammatory granulomas. For immunohistochemistry (IHC) staining, sections were stained with Ly6G (E6Z1T) Rabbit mAb (#87048, Cell Signaling Technology) and counterstained with hematoxylin. H&E staining and IHC staining of slides were stained using the BOND automatic system (Leica) and scanned in Aperio (Leica). The image postprocessing and analysis were done in Aperio ImageScope software (Leica).

For tartrate-resistant acid phosphatase (TRAP) staining, paraffin embedding and tissue sectioning were performed as previously described. Slides were stained according to the TRAP staining kit (#G1050, Servicebio) operating standard, scanned in 3DHISTECH scanner (DANJIER, China), and images were processed in 3DHISTECH SlideViewer software (DANJIER, China).

### Hemogram detection of peripheral blood

Heparinized capillary tubes (#2501, KIMBLE) were accessed to the fundi of mice eyes and rotated to cut the venous plexus. After sufficient blood were drawn, it is transferred to centrifuge tubes with anticoagulant (#110403015, BKMAM). The number and frequency of blood cells were detected by automatic blood cell analyzer (URIT, China).

### Micro-computed tomography

Hind paws were scanned by *in vivo* micro-computed tomography Skyscan 1276 (Bruker). Scanning parameters were voltage, 50 kV; current, 200 μA; filter, 0.25 mm aluminum; pixel size, 9 mm; exposure time, 650 to 800 ms; rotation step, 0.4° for 180° total; object to source distance, 81.719 mm; and camera to source distance, 158.176 mm; with time of scanning, 7 min. Reconstruction of virtual slices was performed in NRecon software 1.7.3 (Bruker) with setup for smoothing = 3, ring artifact correction = 6, and beam hardening correction = 30%. Intensities of interest for reconstruction were in the range of 0.0030 to 0.1500. For reorientation of virtual slices to the same orientation, the DataViewer 1.5.4 software (Bruker) was used. CTvox 3.3.0 (Bruker) was used for Micro-CT image processing.

### ROS detection

To assess ROS production in vivo, mice were intraperitoneal (i.p.) injected with luminescence reporter L-012 (#120-04891, FUJIFILM Wako) in final concentration 25 mg/kg dissolved in PBS with concentration of 20 mmol/L. Luminescence signal was acquired by IVIS Spectrum (PerkinElmer), with the exposure parameters set to automatic. The quantification of photon counts was performed in Living Image Software (PerkinElmer).

### ELISA

Hind paw tissue was cut into small pieces and homogenized with tissue homogenizer (60 HZ, 60 s) in 1 ml RIPA Lysis buffer (#E122-01, Genstar) containing 1 mM PMSF (#B111-01, Genstar). After two rounds of centrifugation (each 12,000 rpm, 10 min, 4°C) the lysates were snap-frozen in liquid nitrogen and stored in −80°C.

Frozen tissue homogenates described above were thawed, total protein concentration was measured by Nanodrop (Thermo). Enzyme linked immunosorbent assay (ELISA) was performed according to manufacturer’s instructions using Mouse IL-1β ELISA Kit (#EK201B, MULTI SCIENCES) and Mouse CCL3/MIP-1a ELISA Kit (#EK261, MULTI SCIENCES) to detect the concentrations of IL-1β and CCL3/MIP-1α in the tissue homogenates.

### Flow cytometry

The bone marrow cells were flushed out with FACS buffer. Single-cell suspensions were treated with red blood cell lysis buffer, stained, and analyzed using BD LSRFortessa (BD Biosciences). For myeloid cell analysis, staining with antibodies against CD11b (#101224, BioLegend) and Gr-1 (#108408, BioLegend). For monocyte and neutrophil analysis, cells were incubated with biotinylated antibodies against c-Kit (#105812, BioLegend), CD11b, Ly-6C (#128024, BioLegend), Ly-6G (#127608, BioLegend). For osteoclast precursor (OCP) analysis, staining with antibodies against c-Kit, CD11b and Csf1r (#135511, BioLegend).

### Real-time quantitative PCR

Hind paws were cut into small pieces and homogenized in 1 mL TRIzol using tissue homogenizer (60 HZ, 60 s). Total RNA was isolated from the hind paws with TRIGene (#P118-05, Genstar) according to the manufacturer’s instructions. In brief, 200 μL of chloroform was added to the 1 mL of TRIGene, and the samples were incubated at room temperate for 5 min following vortexing. After centrifugation, the aqueous phase was transferred to a new DNase/RNase-free tube (BIOFIL), and equal volumes of isopropanol were added. After incubation at room temperature for 10 min, the RNA was pelleted by centrifugation, and then the RNA was washed two times in 75% ethanol before resuspension in DEPC water. 1 μg of RNA was reverse-transcribed to cDNA using StarScript III RT MasterMix (#A233, Genstar) according to the manufacturer’s instructions. Real-time quantitative PCR was performed using a 2×RealStar Power SYBR qPCR Mix (#A314, Genstar) and samples were run on a LightCycler ® 480 Instrument II (Roche). For each sample, transcript levels of tested genes were normalized to Actb.

### Bulk RNA sequencing (RNA-seq) Analysis

For RNA extraction, total RNA was extracted from the tissue using TRIzol^®^ Reagent according the manufacturer’s instructions. Then RNA quality was determined by 5300 Bioanalyser (Agilent) and quantified using the ND-2000 (NanoDrop Technologies). Only high-quality RNA sample (OD260/280=1.8∼2.2, OD260/230≥2.0, RIN≥6.5, 28S:18S≥1.0) was used to construct sequencing library.

For library preparation and sequencing, RNA purification, reverse transcription, library construction and sequencing were performed at Shanghai Majorbio Bio-pharm Biotechnology Co., Ltd. (Shanghai, China) according to the manufacturer’s instructions (Illumina, San Diego, CA). The RNA-seq transcriptome library was prepared following Illumina® Stranded mRNA Prep, Ligation from Illumina (San Diego, CA) using 1μg of total RNA. Shortly, messenger RNA was isolated according to polyA selection method by oligo(dT) beads and then fragmented by fragmentation buffer firstly. Secondly double-stranded cDNA was synthesized using a SuperScript double-stranded cDNA synthesis kit (Invitrogen, CA) with random hexamer primers (Illumina). Then the synthesized cDNA was subjected to end-repair, phosphorylation and ‘A’ base addition according to Illumina’s library construction protocol. Libraries were size selected for cDNA target fragments of 300 bp on 2% low range ultra agarose followed by PCR amplified using Phusion DNA polymerase (NEB) for 15 PCR cycles. After quantified by Qubit 4.0, paired-end RNA-seq sequencing library was sequenced with the NovaSeq 6000 sequencer (2×150 bp read length). For quality control and read mapping, the raw paired end reads were trimmed and quality controlled by fastp with default parameters. Then clean reads were separately aligned to reference genome with orientation mode using HISAT2 software. The mapped reads of each sample were assembled by StringTie in a reference-based approach.

For differential expression analysis and functional enrichment, to identify DEGs between two different samples, the expression level of each transcript was calculated according to the transcripts per million reads (TPM) method. RSEM was used to quantify gene abundances. Essentially, differential expression analysis was performed using the DESeq2 or DEGseq. DEGs with |log2FC|≥1 and FDR≤0.05 (DESeq2) or FDR≤0.001 (DEGseq) were considered to be significantly different expressed genes. In addition, functional-enrichment analysis including GO and KEGG were performed to identify which DEGs were significantly enriched in GO terms and metabolic pathways at Bonferroni-corrected P-value≤0.05 compared with the whole-transcriptome background. GO functional enrichment and KEGG pathway analysis were carried out by Goatools and KOBAS, respectively.

### scRNA-seq and analysis of the scRNA-seq datasets

#### Cell preparation

After harvested, tissues were washed in ice-cold RPMI1640 and dissociated using Collagenase Ⅱ (Sigma 9001-12-1, V900892-100MG) and DNaseⅠ (Sigma 9003-98-9, DN25-1G). Cell count and viability was estimated using fluorescence Cell Analyzer (Countstar^®^ Rigel S2) with AO/PI reagent after removal erythrocytes (Solarbio R1010). Finally, fresh cells were washed twice in the RPMI1640 and then resuspended at 1×10^6^ cells per ml in 1×PBS and 0.04% bovine serum albumin.

### scRNA-seq library construction and sequencing

Single-cell RNA-Seq libraries were prepared using SeekOne^®^ Digital Droplet Single Cell 3’ library preparation kit (SeekGene Catalog No. K00202). Briefly, appropriate number of cells were mixed with reverse transcription reagent and then added to the sample well in SeekOne^®^ chip. Subsequently Barcoded Hydrogel Beads (BHBs) and partitioning oil were dispensed into corresponding wells separately in chip. After emulsion droplet generation reverse transcription were performed at 42 [for 90 minutes and inactivated at 80[for 15 minutes. Next, cDNA was purified from broken droplet and amplified in PCR reaction. The amplified cDNA product was then cleaned, fragmented, end repaired, A-tailed and ligated to sequencing adaptor. Finally, the indexed PCR were performed to amplified the DNA representing 3’ polyA part of expressing genes which also contained Cell Barcode and Unique Molecular Index. The indexed sequencing libraries were cleanup with SPRI beads, analyzed by Qubit (Thermo Fisher Q33226) and Agilent 4200 TapeStation (Agilent G2991BA). The libraries were then sequenced on illumina NovaSeq 6000 with PE150 read length.

### Identifying the marker genes of single cell

The FindAllMarkers function was used to identify marker genes of single cells^[54]^. Aggregation-mediated cell clusters were identified based on DEGs exhibiting log fold changes (logFCs) and aggrephagy-related genes.

### Analysis of transcription factors

SCENIC was employed as a computational tool to simultaneously reconstruct gene regulatory networks and identify stable cell states from single-cell RNA-seq data^[55]^. The inference of the gene regulatory network was based on coexpression analysis and DNA motif analysis, followed by the examination of network activity in individual cells to determine their cellular status^[55, 56]^. We analyzed transcription factors (TFs) using the pySCENIC package^[57]^.

### Cell–cell communication analysis

The signaling inputs and outputs among the different cell types and aggrephagy-mediated cell clusters were assessed using the CellChat package^[58]^. The netVisual_circle function was employed to evaluate the strength of cell[cell communication networks within specific subsets of cells^[58]^.

### Pseudotime analysis

The core concept of CytoTRACE is that cells with a high level of differentiation express fewer genes, while cells with a low level of differentiation express more genes^[59]^. We used the CytoTRACE package to predict the differentiation status of cell subsets.

### Statistical analysis

The *p* values were calculated in GraphPad Prism 9. Comparisions between 2 groups were determined by unpaired t test (two-tailed) for data. Comparison of multiple groups were determined by two-way ANOVA with Tukey-Kramer or Bonferroni multiple comparisons test. Most of experiments in this study were repeated 2 or 3 times independently and representative data were shown. Symbol meanings are as follows: ns, not statistically significant; \**p* < 0.05; \*\**p* < 0.01; \*\*\**p* < 0.001; \*\*\*\**p* < 0.0001. “n” in figure legends represent number of experiment repeats, points in column scatter plots represent biological replicates.

## Conclusion

Although *PSTPIP2* has not been implicated with a direct role in human inflammatory diseases, *Pstpip2^cmo/cmo^* is widely used for testing inflammation regulation and a classic model for CRMO. In the study, we described a new *Pstpip2* frameshift mutations mice strain and confirmed *Pstpip2* loss-of-function causes a CRMO disease in mice. More importantly, we demonstrated that loss of *Morrbid* through various layers of mechanisms inhibited the occurrence of the disease (Figure 6). Our results not only demonstrated the link between *Morrbid* and AIDs and the function of *Morrbid* in a new special type of myeloid cells, osteoclasts, but also conducted single-cell transcriptome sequencing technologies to reveal the CRMO pathogenesis induced by loss of *Pstpip2* and the intervention by the loss of *Morrbid*. In conclusion, our study describes that CRMO occurrence relies on multiple layers of inflammatory myeloid cells at the single-cell level and provides a theoretical and preclinical basis for *Morrbid* as a therapeutic target for autoinflammatory diseases.

**Figure 6.**
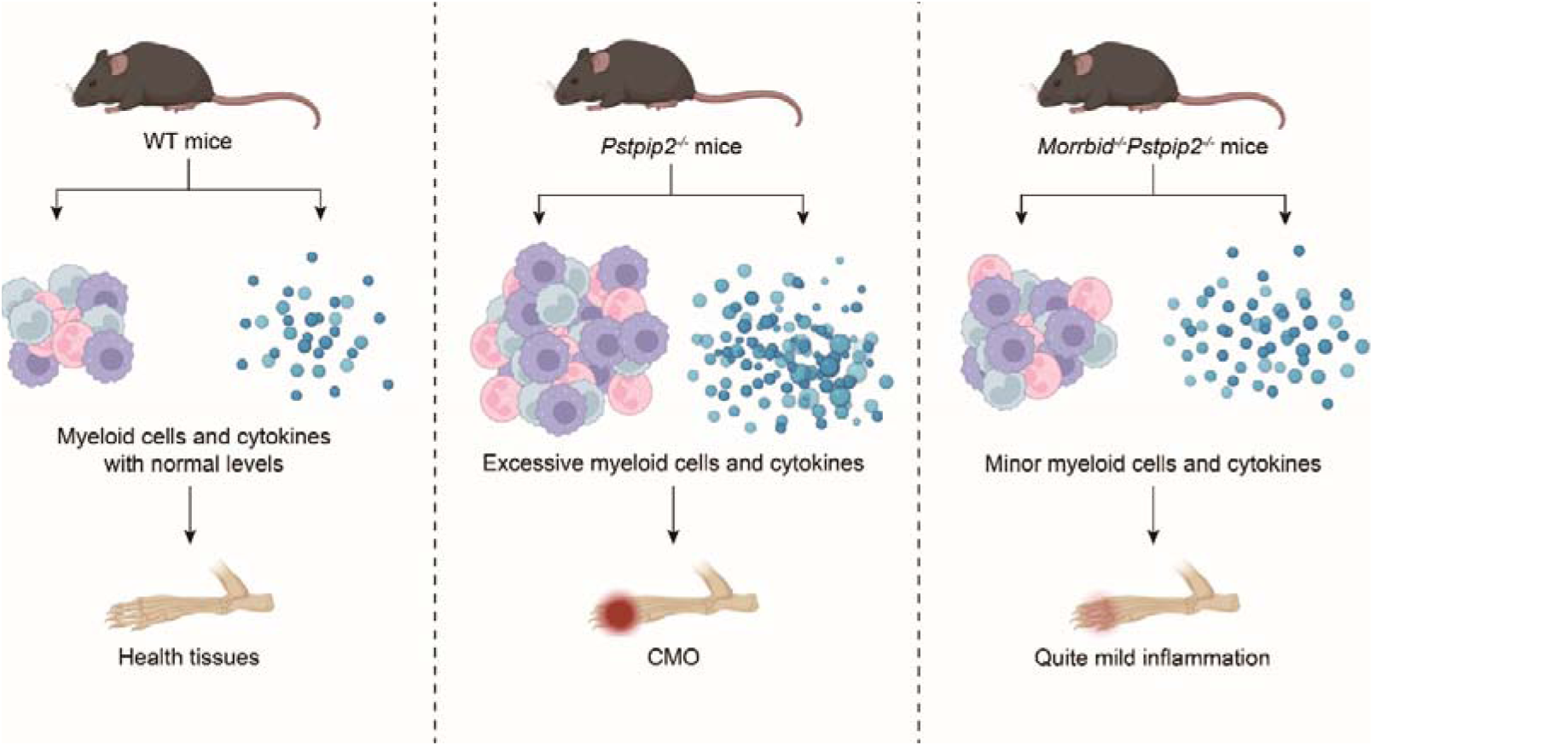
Mouse models with genetic mutation in *Pstpip2* are one of the most important and fantastic models for studying autoinflammatory diseases (AIDs) including osteomyelitis. We generated a 5-bp deletion in *Pstpip2* for testing *Morrbid* involvement in the AIDs. Our genetic analysis and pathological analysis including scRNA-seq profiling demonstrate that *Morrbid* is an effective target for anti-inflammation and loss of *Morrbid* mitigate osteomyelitis induced by loss of *Pstpip2*.

## Data availability

Any additional information required to reanalyze the data reported in this paper is available from the lead contact upon request.

The raw sequencing data from this study have been deposited in the Genome Sequence Archive in BIG Data Center (https://bigd.big.ac.cn/), Beijing Institute of Genomics (BIG), Chinese Academy of Sciences, under the accession number: PRJCA026323.

## Authors’ contributions and materials

ZC and QH conceived and designed the study, drafted and wrote the manuscript. ZC and JD wrote the computational scripts for sequencing analysis and visualized the results. QH performed most of the experiments and analysis.

## Funding

This work was supported by grants from Tianjin Municipal Education Commission Scientific Research Program Projects (#2022KJ194 to DG).

## Acknowledgments

We thank members of Cai laboratory and colleagues of Tianjin Medical University for their technical and administration support, as well as for their helpful suggestions improve the manuscript.

## Declarations section

### Competing interests

ZC is a scientific advisor to Beijing SeekGene BioSciences Co. Ltd. Other authors declare no potential conflict of interest.

---

**Supplementary Figure 1.**
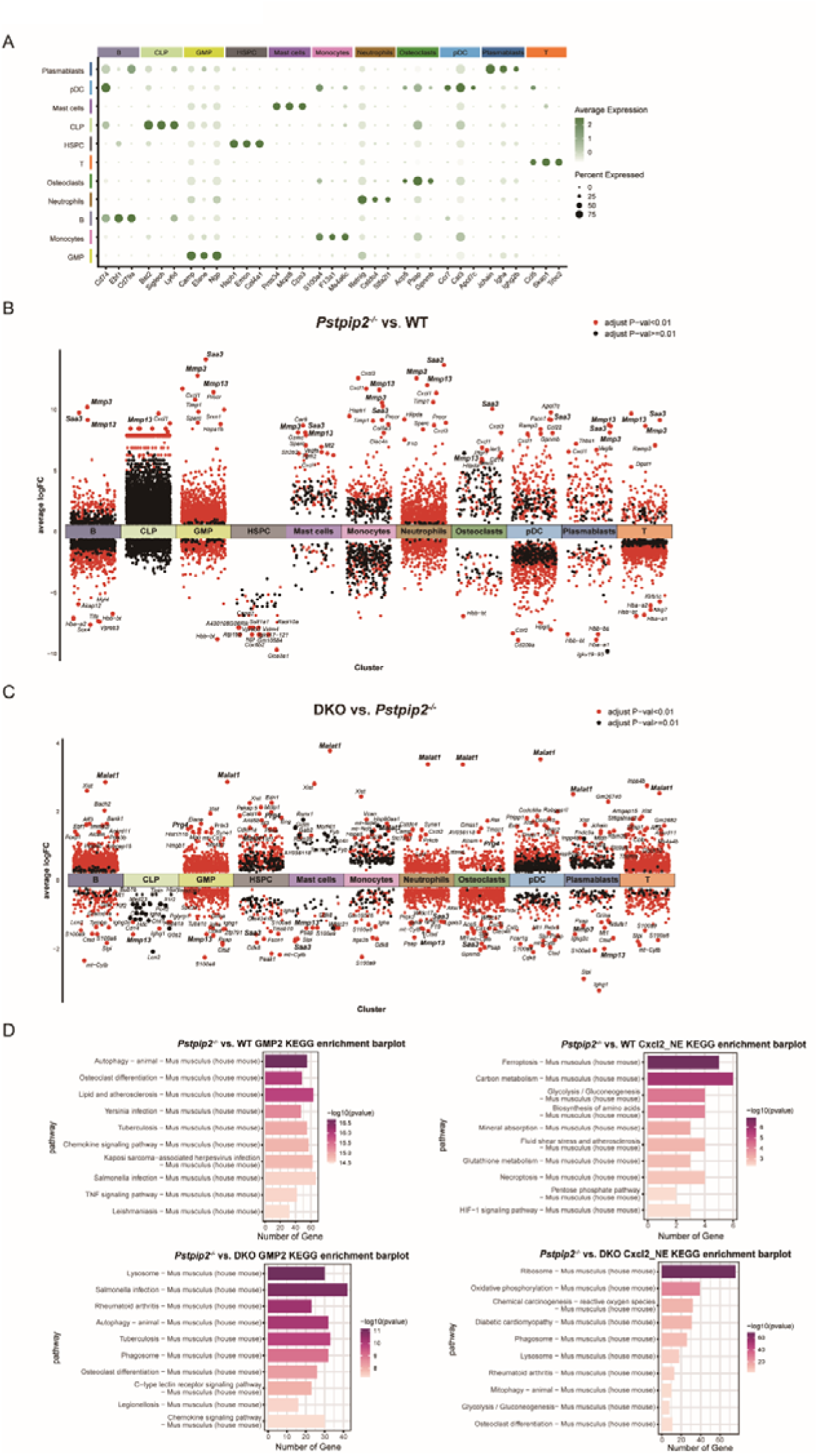
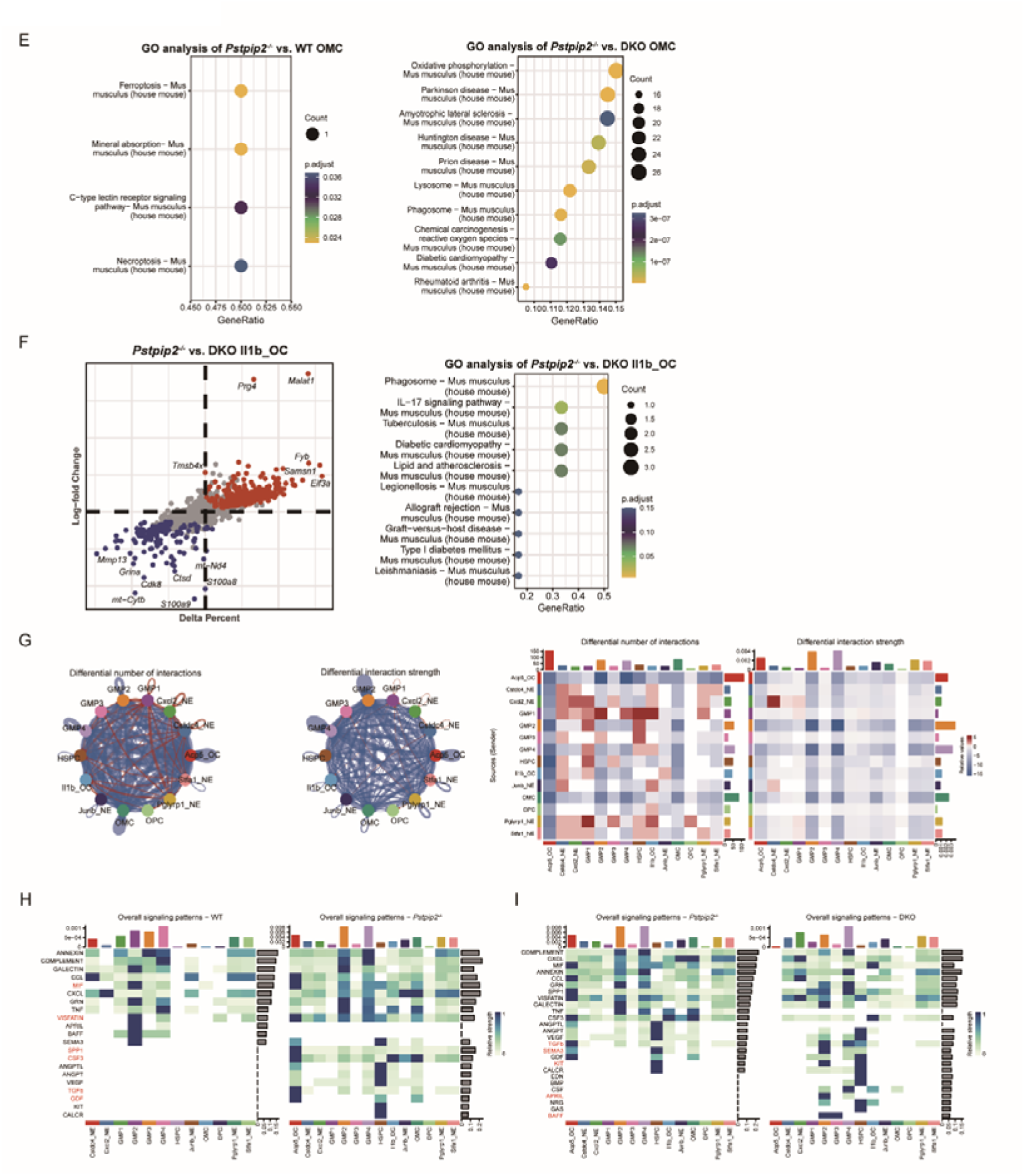
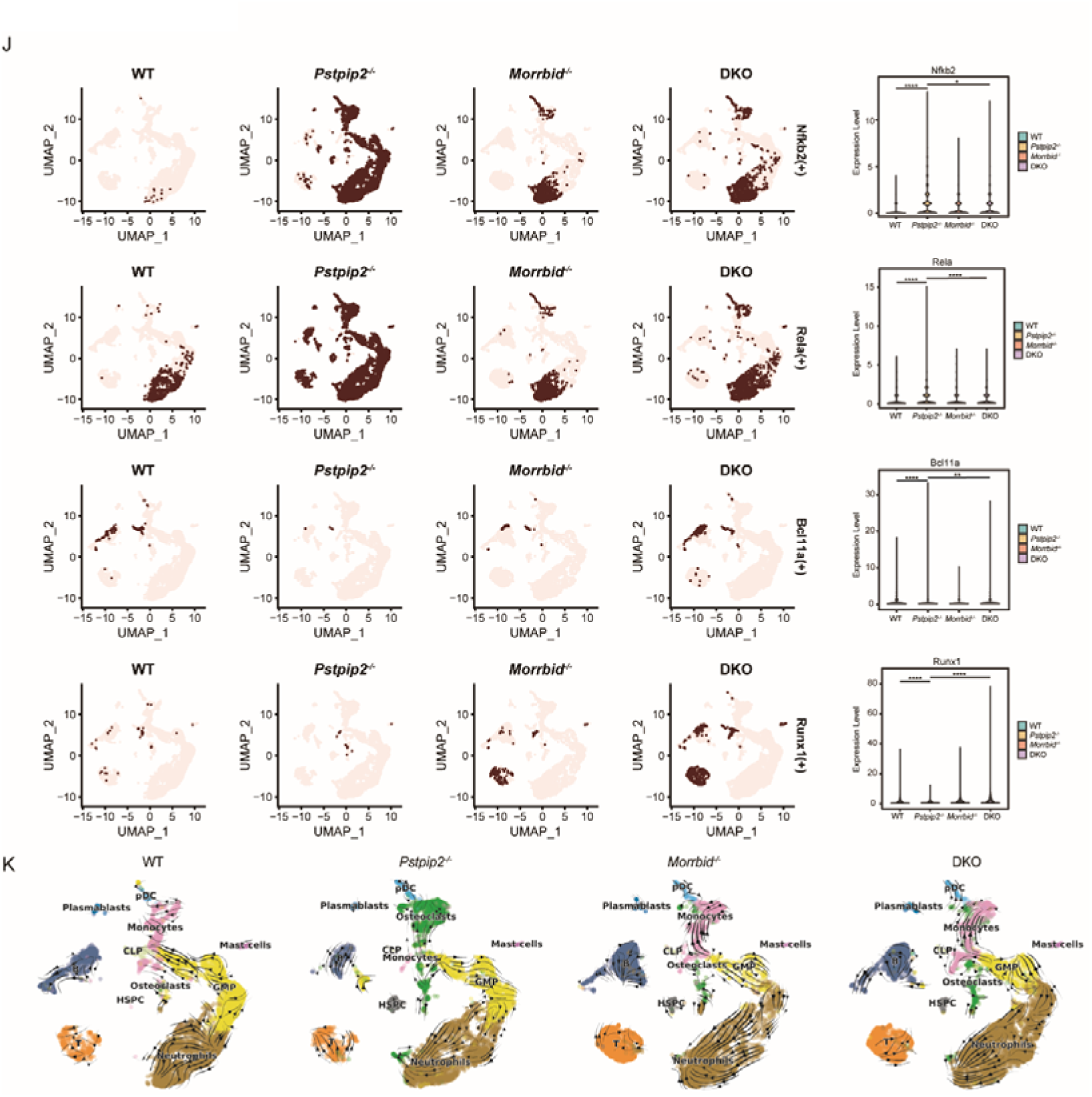
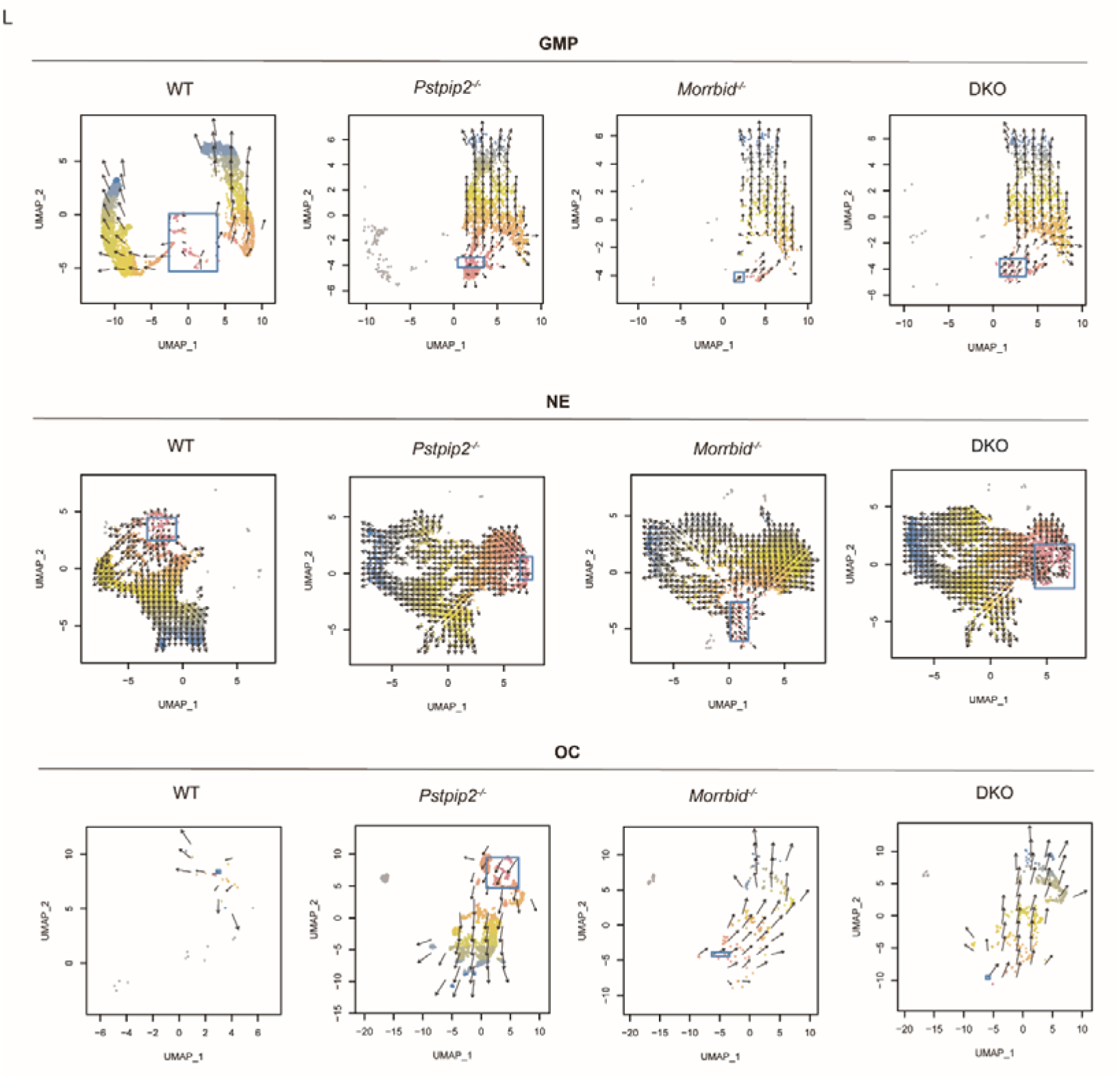
Supplementary Figure Additional computational analysis of scRNA-seq datasets from the four genotype mice. (**A**) Expression profiles of marker genes used for annotating the BM cells in the scRNA-seq datasets. (**B** and **C**) Volcano plots showing DEGs in each cell-type compartment from *Pstpip2^−/−^*mice compared with that from WT and DKO mice. (**D**) KEGG enrichment analysis of GMP2 and Cxcl2_NE among different groups. (**E**) GO enrichment analysis of osteoclast/monocytes (OMC) among different groups. (**F**) Scatter plots showing DEGs in the indicated comparison. (**G**) Circle plot (left) and heatmap (right) of the relative number and strength of interactions in DKO compared with *Pstpip2^−/−^* mice; the up-regulated intercellular interactions are shown in red; the down-regulated intercellular interactions are represented in blue. (**H** and **I**) Heatmap for overall signal patterns of WT compared with *Pstpip2^−/−^* mice; and for *Pstpip2^−/−^* mice compared with DKO mice. (**J**) Expression (left) and quantification (right) of representative transcription factors in the four genotype mice. Transcriptional factors Nfkb2, Rela, Bcl11a and Runx1 are included. (**K**) RNA velocity in all BM cells from the four groups of mice. (**L**) Developmental trajectories of GMP, NE and OC in four groups of mice. Developmental initiation points are marked by blue boxes.

**Supplementary Table.**
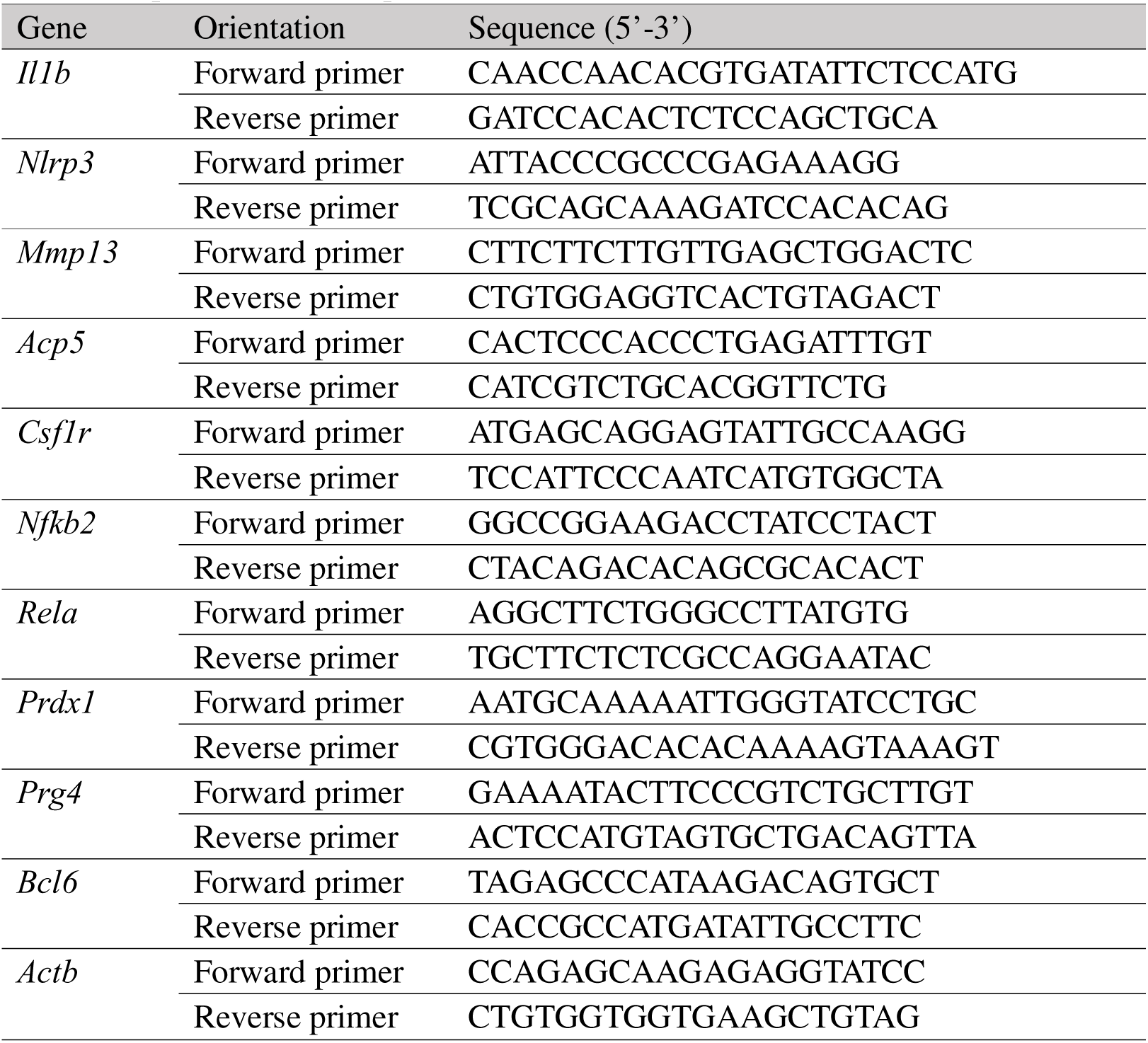
Primer sequences used for quantitative RT-PCR are listed below:

## Notes

### Competing Interest Statement

The authors have declared no competing interest.

## References

[1] Krainer, J., Siebenhandl, S., Weinhäusel, A., Systemic autoinflammatory diseases, J Autoimmun 109 (2020) 102421.

[2] Szekanecz, Z., McInnes, I. B., Schett, G., et al., Autoinflammation and autoimmunity across rheumatic and musculoskeletal diseases, Nat Rev Rheumatol 17 (2021) 585–595.

[3] Meier-Schiesser, B., French, L. E., Autoinflammatory syndromes, J Dtsch Dermatol Ges 19 (2021) 400–426.

[4] Rood, J. E., Behrens, E. M., Inherited Autoinflammatory Syndromes, Annu Rev Pathol 17 (2022) 227–249.

[5] Beck, D. B., Ferrada, M. A., Sikora, K. A., et al., Somatic Mutations in UBA1 and Severe Adult-Onset Autoinflammatory Disease, New England Journal of Medicine 383 (2020) 2628–2638.

[6] Georgin-Lavialle, S., Fayand, A., Rodrigues, F., et al., Autoinflammatory diseases: State of the art, Presse Med 48 (2019) e25–e48.

[7] Stern, S. M., Ferguson, P. J., Autoinflammatory bone diseases, Rheum Dis Clin North Am 39 (2013) 735–49.

[8] Cox, A. J., Ferguson, P. J., Update on the genetics of nonbacterial osteomyelitis in humans, Curr Opin Rheumatol 30 (2018) 521–525.

[9] Cox, A. J., Darbro, B. W., Laxer, R. M., et al., Recessive coding and regulatory mutations in FBLIM1 underlie the pathogenesis of chronic recurrent multifocal osteomyelitis (CRMO), PLoS One 12 (2017) e0169687.

[10] Wang, Y., Wang, J., Zheng, W., et al., Identification of an IL-1 receptor mutation driving autoinflammation directs IL-1-targeted drug design, Immunity 56 (2023) 1485–1501.e7.

[11] Chitu, V., Nacu, V., Charles, J. F., et al., PSTPIP2 deficiency in mice causes osteopenia and increased differentiation of multipotent myeloid precursors into osteoclasts, Blood 120 (2012) 3126–35.

[12] Cassel, S. L., Janczy, J. R., Bing, X., et al., Inflammasome-independent IL-1β mediates autoinflammatory disease in Pstpip2-deficient mice, Proc Natl Acad Sci U S A 111 (2014) 1072–7.

[13] Yao, Y., Cai, X., Zhang, M., et al., PSTPIP2 regulates synovial macrophages polarization and dynamics via ERβ in the joint microenvironment, Arthritis Res Ther 24 (2022) 247.

[14] Kralova, J., Drobek, A., Prochazka, J., et al., Dysregulated NADPH Oxidase Promotes Bone Damage in Murine Model of Autoinflammatory Osteomyelitis, J Immunol 204 (2020) 1607–1620.

[15] Xu, J. J., Li, H. D., Du, X. S., et al., Role of the F-BAR Family Member PSTPIP2 in Autoinflammatory Diseases, Front Immunol 12 (2021) 585412.

[16] Kralova, J., Pavliuchenko, N., Fabisik, M., et al., The receptor-type protein tyrosine phosphatase CD45 promotes onset and severity of IL-1β-mediated autoinflammatory osteomyelitis, J Biol Chem 297 (2021) 101131.

[17] Lukens, J. R., Gurung, P., Vogel, P., et al., Dietary modulation of the microbiome affects autoinflammatory disease, Nature 516 (2014) 246–9.

[18] Zhang, D., Su, G., Liu, Y., et al., Clinical and genetic characteristics of PSTPIP1-associated myeloid-related proteinemia inflammatory syndrome, Pediatr Rheumatol Online J 19 (2021) 151.

[19] Kotzin, J. J., Spencer, S. P., McCright, S. J., et al., The long non-coding RNA Morrbid regulates Bim and short-lived myeloid cell lifespan, Nature 537 (2016) 239–243.

[20] Cai, Z., Kotzin, J. J., Ramdas, B., et al., Inhibition of Inflammatory Signaling in Tet2 Mutant Preleukemic Cells Mitigates Stress-Induced Abnormalities and Clonal Hematopoiesis, Cell Stem Cell 23 (2018) 833–849.e5.

[21] Cai, Z., Lu, X., Zhang, C., et al., Hyperglycemia cooperates with Tet2 heterozygosity to induce leukemia driven by proinflammatory cytokine-induced lncRNA Morrbid, J Clin Invest 131 (2021)

[22] Kotzin, J. J., Iseka, F., Wright, J., et al., The long noncoding RNA Morrbid regulates CD8 T cells in response to viral infection, Proc Natl Acad Sci U S A 116 (2019) 11916–11925.

[23] Zohar, K., Lezmi, E., Reichert, F., et al., Coordinated Transcriptional Waves Define the Inflammatory Response of Primary Microglial Culture, Int J Mol Sci 24 (2023)

[24] Baslan, T., Hicks, J., Unravelling biology and shifting paradigms in cancer with single-cell sequencing, Nature Reviews Cancer 17 (2017) 557–569.

[25] Yokota, K., Sato, K., Miyazaki, T., et al., Characterization and Function of Tumor Necrosis Factor and Interleukin-6-Induced Osteoclasts in Rheumatoid Arthritis, Arthritis Rheumatol 73 (2021) 1145–1154.

[26] Yao, Z., Getting, S. J., Locke, I. C., Regulation of TNF-Induced Osteoclast Differentiation, Cells 11 (2021)

[27] Mori, T., Miyamoto, T., Yoshida, H., et al., IL-1β and TNFα-initiated IL-6-STAT3 pathway is critical in mediating inflammatory cytokines and RANKL expression in inflammatory arthritis, Int Immunol 23 (2011) 701–12.

[28] Roodman, G. D., Biology of osteoclast activation in cancer, J Clin Oncol 19 (2001) 3562–71.

[29] Toh, K., Kukita, T., Wu, Z., et al., Possible involvement of MIP-1alpha in the recruitment of osteoclast progenitors to the distal tibia in rats with adjuvant-induced arthritis, Lab Invest 84 (2004) 1092–102.

[30] Roodman, G. D., Regulation of osteoclast differentiation, Ann N Y Acad Sci 1068 (2006) 100–9.

[31] Griffith, B., Pendyala, S., Hecker, L., et al., NOX enzymes and pulmonary disease, Antioxid Redox Signal 11 (2009) 2505–16.

[32] Mittal, M., Siddiqui, M. R., Tran, K., et al., Reactive oxygen species in inflammation and tissue injury, Antioxid Redox Signal 20 (2014) 1126–67.

[33] Agidigbi, T. S., Kim, C., Reactive Oxygen Species in Osteoclast Differentiation and Possible Pharmaceutical Targets of ROS-Mediated Osteoclast Diseases, Int J Mol Sci 20 (2019)

[34] Jarczak, D., Nierhaus, A., Cytokine Storm-Definition, Causes, and Implications, Int J Mol Sci 23 (2022)

[35] Ono, T., Nakashima, T., Recent advances in osteoclast biology, Histochem Cell Biol 149 (2018) 325–341.

[36] Hayman, A. R., Tartrate-resistant acid phosphatase (TRAP) and the osteoclast/immune cell dichotomy, Autoimmunity 41 (2008) 218–23.

[37] Ballanti, P., Minisola, S., Pacitti, M. T., et al., Tartrate-resistant acid phosphate activity as osteoclastic marker: sensitivity of cytochemical assessment and serum assay in comparison with standardized osteoclast histomorphometry, Osteoporos Int 7 (1997) 39–43.

[38] Halling Linder, C., Ek-Rylander, B., Krumpel, M., et al., Bone Alkaline Phosphatase and Tartrate-Resistant Acid Phosphatase: Potential Co-regulators of Bone Mineralization, Calcif Tissue Int 101 (2017) 92–101.

[39] Harada, K., Itoh, H., Kawazoe, Y., et al., Polyphosphate-mediated inhibition of tartrate-resistant acid phosphatase and suppression of bone resorption of osteoclasts, PLoS One 8 (2013) e78612.

[40] de Almeida, L. G. N., Thode, H., Eslambolchi, Y., et al., Matrix Metalloproteinases: From Molecular Mechanisms to Physiology, Pathophysiology, and Pharmacology, Pharmacol Rev 74 (2022) 712–768.

[41] Ji, B., Ma, Y., Wang, H., et al., Activation of the P38/CREB/MMP13 axis is associated with osteoarthritis, Drug Des Devel Ther 13 (2019) 2195–2204.

[42] Hu, Q., Ecker, M., Overview of MMP-13 as a Promising Target for the Treatment of Osteoarthritis, Int J Mol Sci 22 (2021)

[43] Vallon, R., Freuler, F., Desta-Tsedu, N., et al., Serum amyloid A (apoSAA) expression is up-regulated in rheumatoid arthritis and induces transcription of matrix metalloproteinases, J Immunol 166 (2001) 2801–7.

[44] Lee, J. Y., Hall, J. A., Kroehling, L., et al., Serum Amyloid A Proteins Induce Pathogenic Th17 Cells and Promote Inflammatory Disease, Cell 180 (2020) 79–91.e16.

[45] Yao, Q., Wu, X., Tao, C., et al., Osteoarthritis: pathogenic signaling pathways and therapeutic targets, Signal Transduct Target Ther 8 (2023) 56.

[46] Zhao, Y., Ning, J., Teng, H., et al., Long noncoding RNA Malat1 protects against osteoporosis and bone metastasis, Nat Commun 15 (2024) 2384.

[47] Huang, X., Jie, S., Li, W., et al., GATA4-activated lncRNA MALAT1 promotes osteogenic differentiation through inhibiting NEDD4-mediated RUNX1 degradation, Cell Death Discov 9 (2023) 150.

[48] Alnajjar, F. A., Sharma-Oates, A., Wijesinghe, S. N., et al., The Expression and Function of Metastases Associated Lung Adenocarcinoma Transcript-1 Long Non-Coding RNA in Subchondral Bone and Osteoblasts from Patients with Osteoarthritis, Cells 10 (2021)

[49] Massengale, M., Massengale, J. L., Benson, C. R., et al., Adult Prg4+ progenitors repair long-term articular cartilage wounds in vivo, JCI Insight 8 (2023)

[50] Hofmann, S. R., Kapplusch, F., Girschick, H. J., et al., Chronic Recurrent Multifocal Osteomyelitis (CRMO): Presentation, Pathogenesis, and Treatment, Curr Osteoporos Rep 15 (2017) 542–554.

[51] Ferguson, P. J., Laxer, R. M., New discoveries in CRMO: IL-1β, the neutrophil, and the microbiome implicated in disease pathogenesis in Pstpip2-deficient mice, Semin Immunopathol 37 (2015) 407–12.

[52] Chitu, V., Ferguson, P. J., de Bruijn, R., et al., Primed innate immunity leads to autoinflammatory disease in PSTPIP2-deficient cmo mice, Blood 114 (2009) 2497–505.

[53] Snider, C. E., Wan Mohamad Noor, W. N. I., Nguyen, N. T. H., et al., The state of F-BAR domains as membrane-bound oligomeric platforms, Trends Cell Biol 31 (2021) 644–655.

[54] Slovin, S., Carissimo, A., Panariello, F., et al., Single-Cell RNA Sequencing Analysis: A Step-by-Step Overview, Methods Mol Biol 2284 (2021) 343–365.

[55] Kumar, N., Mishra, B., Athar, M., et al., Inference of Gene Regulatory Network from Single-Cell Transcriptomic Data Using pySCENIC, Methods Mol Biol 2328 (2021) 171–182.

[56] Van de Sande, B., Flerin, C., Davie, K., et al., A scalable SCENIC workflow for single-cell gene regulatory network analysis, Nat Protoc 15 (2020) 2247–2276.

[57] Schmitt, P., Sorin, B., Frouté, T., et al., GReNaDIne: A Data-Driven Python Library to Infer Gene Regulatory Networks from Gene Expression Data, Genes (Basel) 14 (2023)

[58] Jin, S., Guerrero-Juarez, C. F., Zhang, L., et al., Inference and analysis of cell-cell communication using CellChat, Nat Commun 12 (2021) 1088.

[59] Gulati, G. S., Sikandar, S. S., Wesche, D. J., et al., Single-cell transcriptional diversity is a hallmark of developmental potential, Science 367 (2020) 405–411.

