## Supplementary figures and images for "Depicting pathogenesis of osteomyelitis by single cell RNA-sequencing and an involvement of *Morrbid* in the autoinflammatory disease"

### Supplementary Figure 1

#### Supplementary Figure

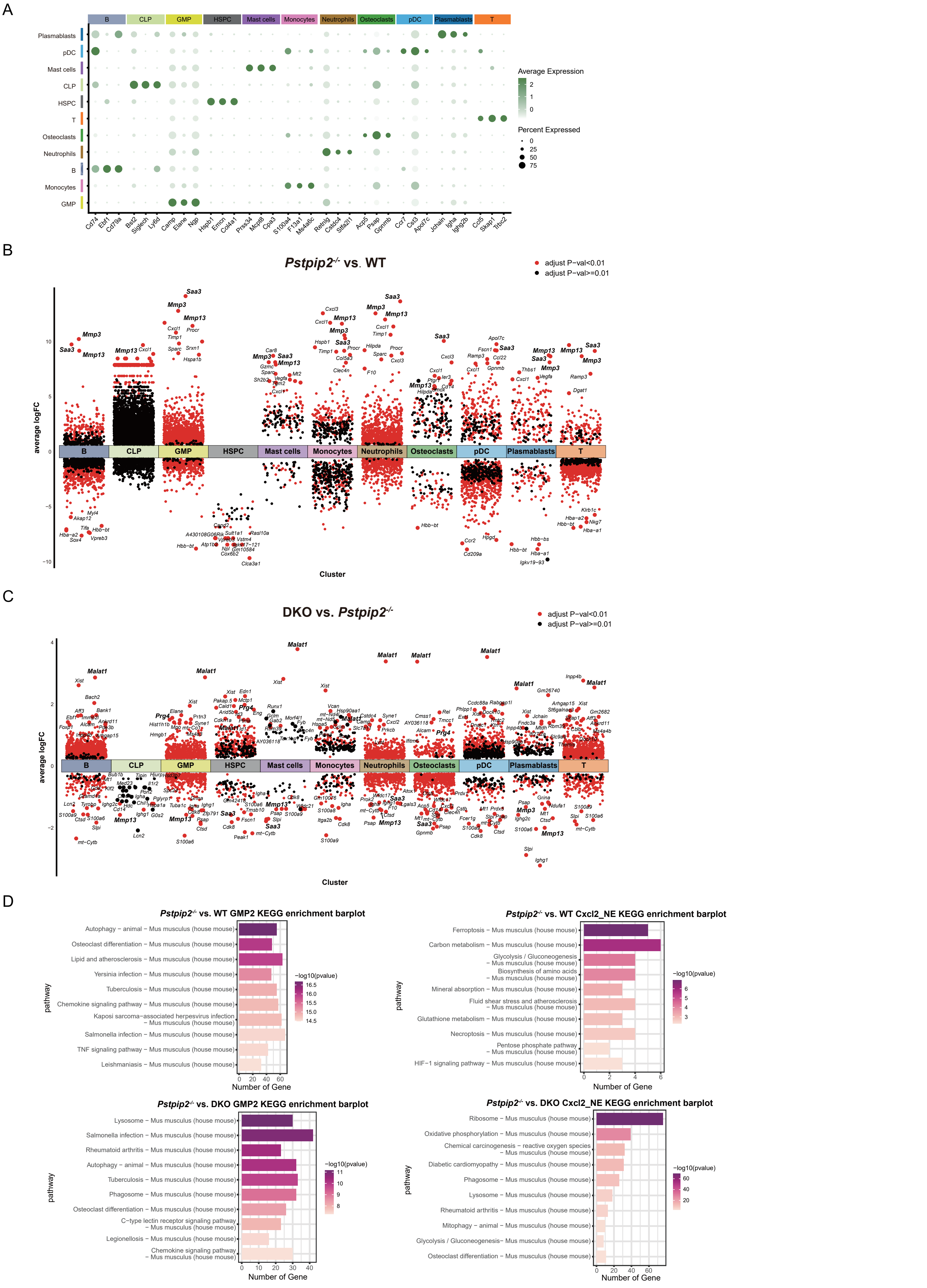

### Supplementary Figure 1

#### Supplementary Figure

E

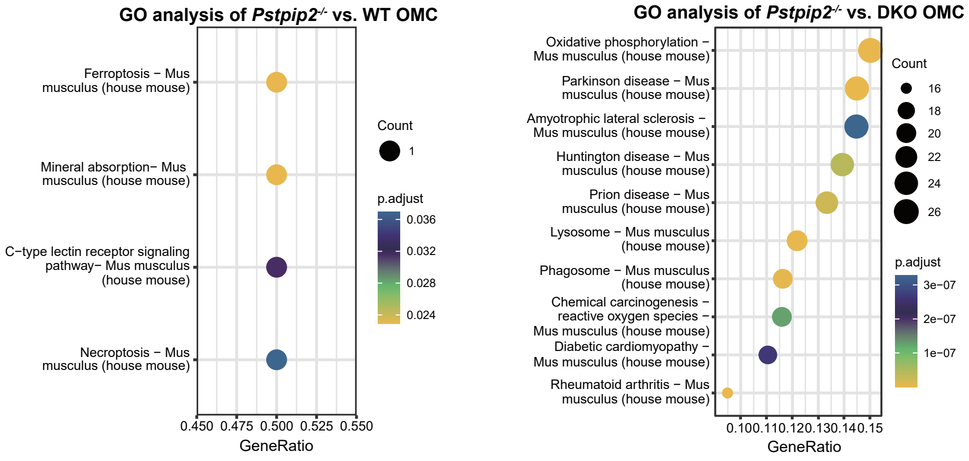


F

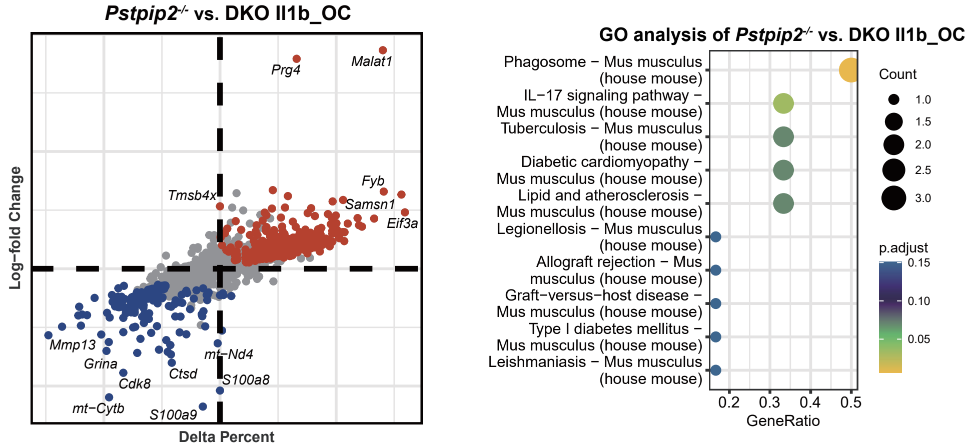


G

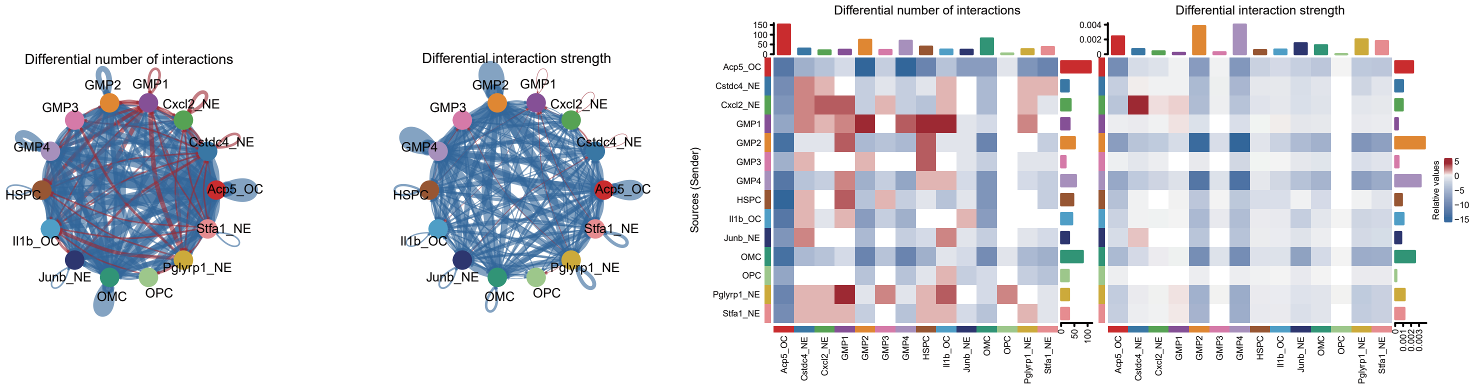


H

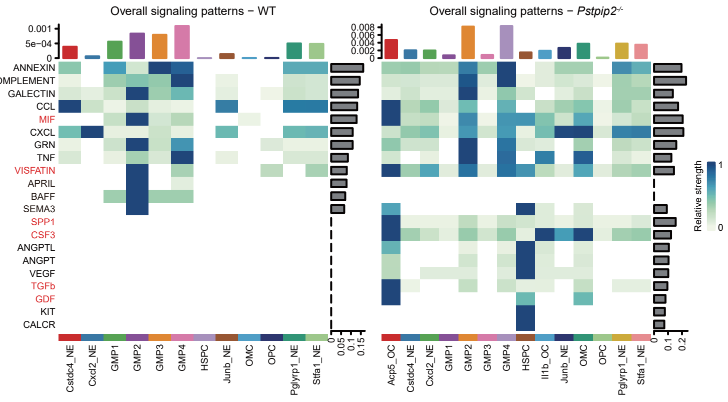


I

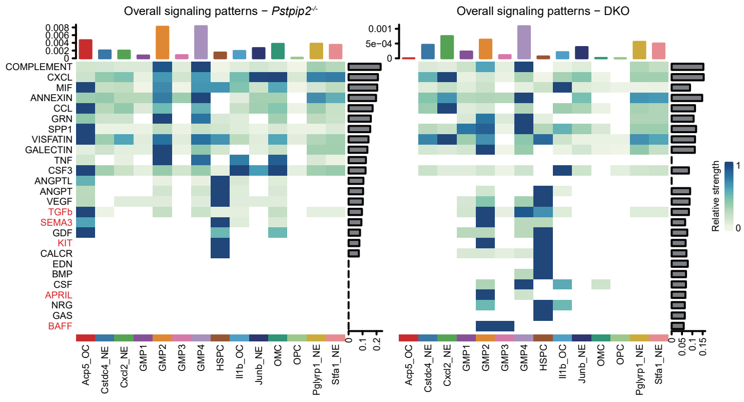

### Supplementary Figure 1

#### Supplementary Figure

**J**

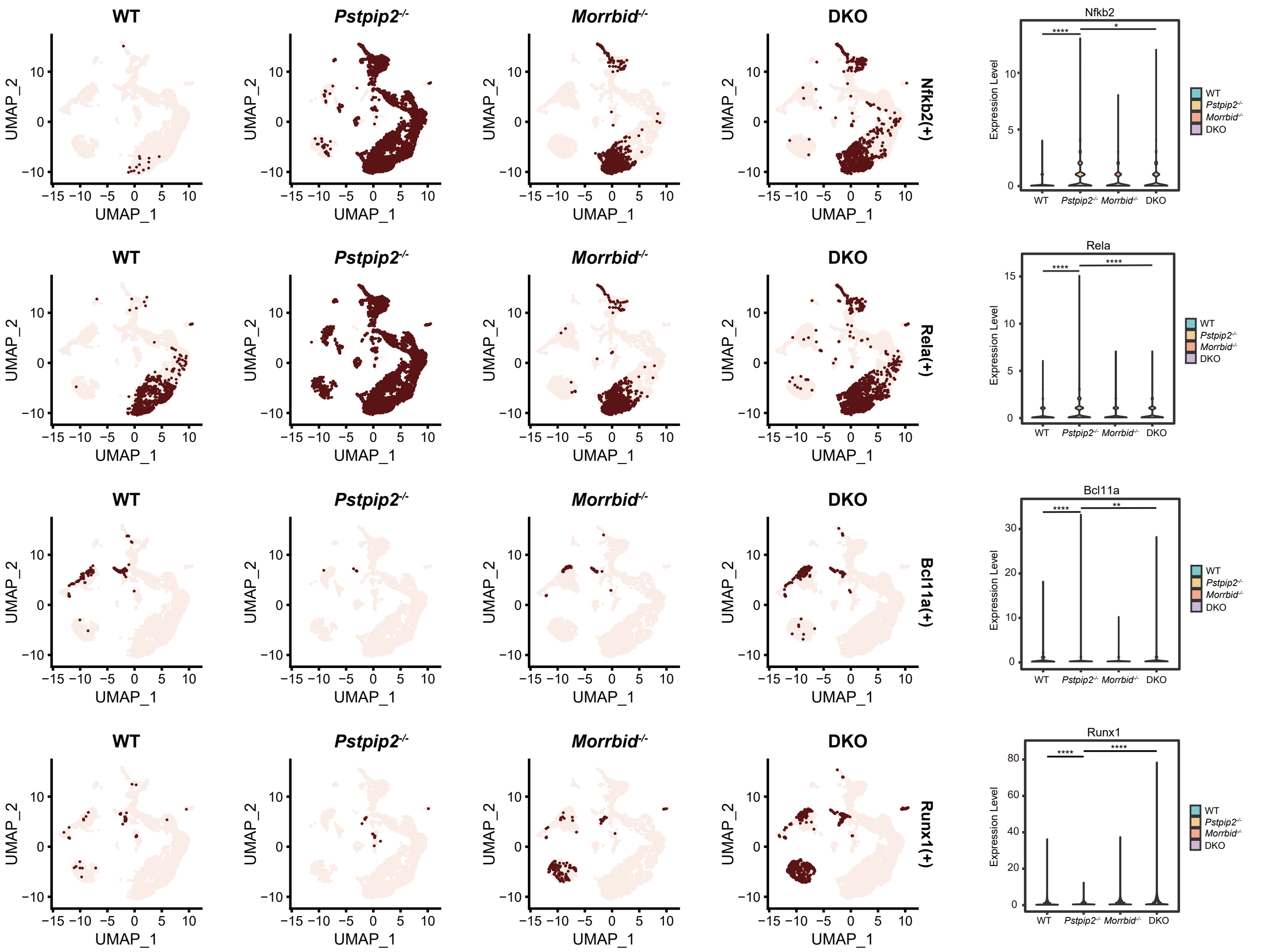


**K**

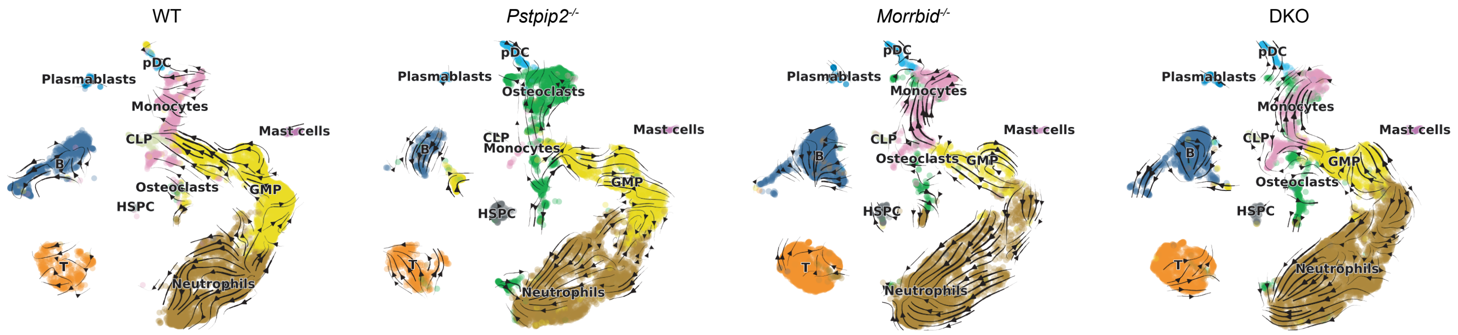

### Supplementary Figure 1

##### Supplementary Figure

**L**

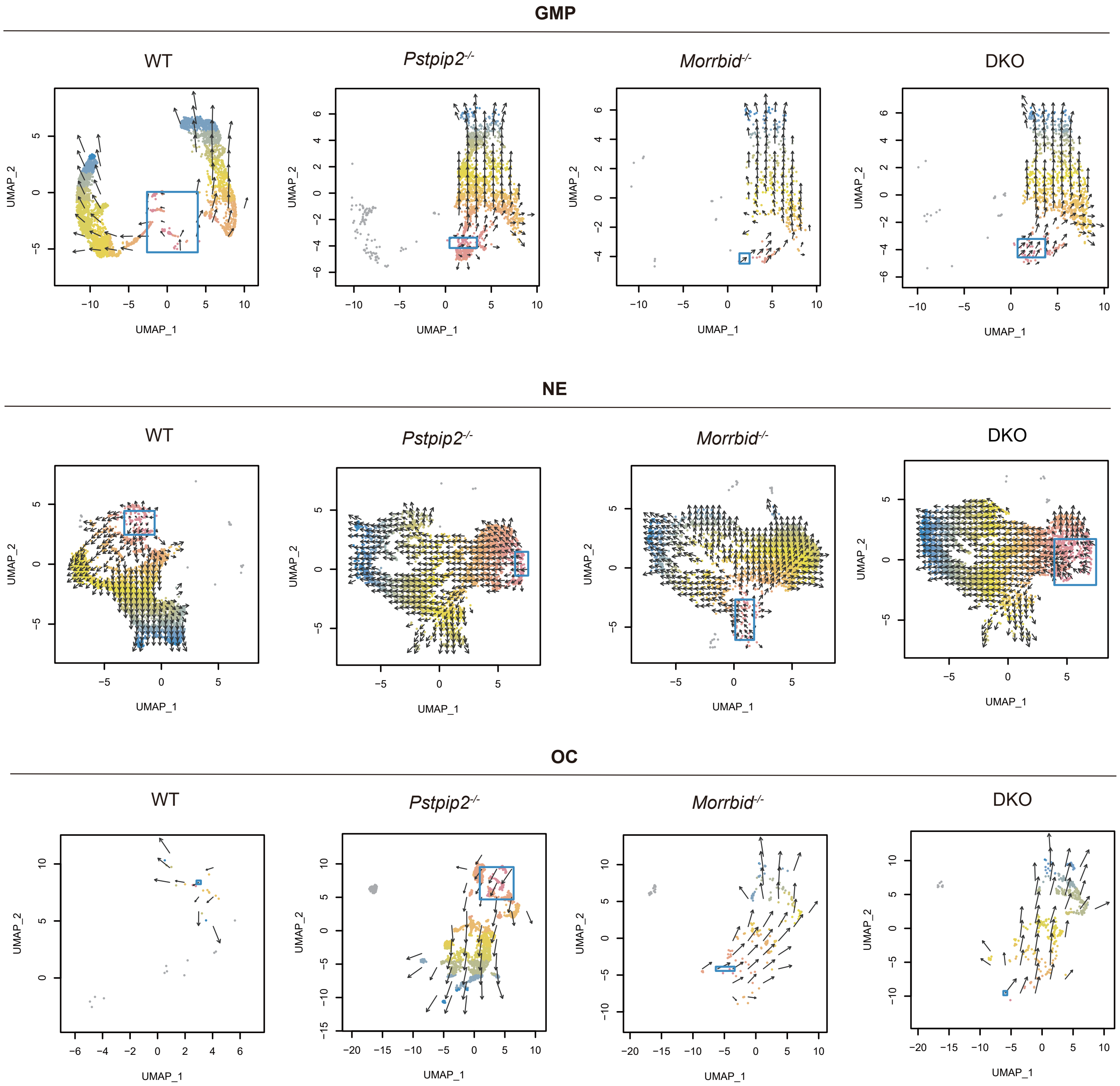
